# Vulnerability segmentation across the fine root-to-leaf hydraulic pathway: Contrasting embolism resistance and dehydration dynamics in eight temperate tree species

**DOI:** 10.64898/2026.11.27.769719

**Authors:** Franziska Geuchen, Christoph Leuschner, Leonie von Rudorff, Sharath S. Paligi

## Abstract

Vulnerability segmentation (VS), whereby distal organs embolize at less negative water potentials than proximal organs, may contribute to drought tolerance, yet its variation across species and relationship with isohydricity remain poorly understood.
We characterized VS across eight temperate tree species using optical vulnerability measurements of leaves/twigs, stems, and coarse and fine roots, quantified embolism thresholds (P_12_, P_50_, P_88_) and dehydration times (T_12_, T_50_, T_88_), and tested their relationships with species minimum epidermal conductance. Embolism thresholds and dehydration time varied substantially among organs, with stems typically showing longer dehydration times than leaves and roots (especially at P_88_).
Four of eight species showed significant segmentation between leaves/fine roots and stems in both embolism thresholds and dehydration time, unrelated to species isohydricity. More negative embolism thresholds did not consistently translate into longer times to hydraulic failure, suggesting decoupling of embolism resistance and dehydration kinetics. Shorter dehydration times were associated with higher minimum conductance, consistent with greater residual water loss.
Vulnerability segmentation is common, yet species-specific; progression to hydraulic failure depends on organ identity and minimum conductance. Thus, embolism resistance and dehydration resistance are complementary, but independent, dimensions of drought response and accurate assessment of drought strategies requires integration of both across plant compartments.

**Highlight:** Xylem vulnerability is species- and organ specific: water potential thresholds and dehydration time to critical embolism levels are independent of species’ isohydricity, underscoring the need to integrate temporal dynamics across whole-plant.

## Introduction

Climate change, particularly the increasing frequency of hotter and severe droughts, poses a major threat to forest ecosystems, accelerating large-scale tree mortality (Allen et al. 2010; Brodribb 2020; Hammond et al. 2022; Pörtner et al. 2022). The main mechanisms of drought-induced tree mortality are hydraulic failure and carbon starvation (McDowell et al. 2008). Hydraulic failure occurs when drought-induced embolisms (gas bubbles) form and spread in the xylem through inter-conduit pit membranes (Lens et al. 2022), blocking water transport (Choat et al. 2018).

Xylem embolism resistance is commonly assessed by xylem vulnerability curves (Sperry and Tyree 1988; Tyree 2003) and is one of the key indicators of species’ drought vulnerability (Anderegg et al. 2016). Xylem embolism resistance is commonly quantified through P_50_, the water potential (Ψ) causing a 50% loss of hydraulic conductance (Choat et al. 2012). P_50_ was identified as the point of no return in gymnosperm species (Urli et al. 2013). Other important thresholds include P_12_, indicating embolism onset when stomates are mostly closed and turgor is lost (Creek et al. 2020; Sorek et al. 2022; Paligi et al. 2024). P_88_ has been linked with catastrophic hydraulic failure and mortality in angiosperm species (Urli et al. 2013; Brodribb and Cochard 2009). Although these thresholds provide useful benchmarks, recent evidence suggests species-specific variation in the point of no return (Hammond et al. 2019; Arend et al. 2021; Mantova et al. 2021) underscoring the need to consider species-specific thresholds.

Drought induced tree mortality is thought to be influenced by species’ isohydric and anisohydric behaviour – the degree to which a species regulates leaf water potential through stomatal closure (Martínez-Vilalta and Garcia-Forner 2017). Isohydric species maintain tighter control, closing stomates early, allowing the leaves to maintain relatively constant water potential throughout the day (small hydroscape area, HSA; Meinzer et al. 2016), but this also limits photosynthesis and risks carbon starvation, which may lead to increased susceptibility to biotic stressors (McDowell et al. 2008). In contrast, anisohydric species tend to keep stomates open longer, maintaining photosynthetic functioning, but with decreasing water potential (greater HSA) risking hydraulic failure (McDowell et al. 2008; Meinzer et al. 2016). However, recent studies show that the timing of stomatal closure and plant dry-down rates relate to the dehydration tolerance of tissues rather than to the degree of isohydricity (Martínez-Vilalta and Garcia-Forner 2017; Blackman et al. 2019; Leuschner et al. 2019; Arend et al. 2021; Paligi et al. 2024, 2026). Thus, embolism timing can provide important insights into species-specific drought responses and the progression of hydraulic failure and may be a key factor in predicting a species’ overall drought vulnerability (Blackman et al. 2016; Martin-StPaul et al. 2017; Duursma et al. 2019). Embolism resistance thresholds can vary within a plant, a phenomenon known as vulnerability segmentation (VS) – a strategy to protect more permanent parts of the plant such as stems and coarse roots by sacrificing temporary organs like leaves, needles, and fine roots (Zimmermann 1983; Tyree and Ewers 1991). However, not all species follow this pattern; for example, some gymnosperm species showed higher embolism resistance in branches than in stems (McCulloh et al. 2014; Zambonini et al. 2024). Because most studies measure xylem embolism resistance in branches (e.g. Kahmen et al. 2022; Schumann et al. 2024), this can lead to under-or overestimation of a species’ drought resistance. Evidence in support of VS varies: some studies report clear segmentation (Hochberg et al. 2017; Rodriguez-Dominguez et al. 2018; Wason et al. 2018; Song et al. 2022; Harrison Day et al. 2024; Haverroth et al. 2024;), while others find little or none (Wason et al. 2018; Wu et al. 2020; Lübbe et al. 2022), suggesting VS is likely species-specific (Cardoso et al. 2022). Hartmann et al. (2021) proposed that VS might be related to the degree of isohydricity, with more isohydric species exhibiting greater VS than anisohydric taxa (Wason et al. 2018; Losso et al. 2019). However, the relationship between VS and isohydric behaviour has not been tested and warrants further investigation.

The optical method enables non-invasive monitoring of embolism formation across multiple plant organs simultaneously (Brodribb et al. 2016; Brodribb et al. 2017). By capturing both the temporal and spatial patterns of embolism formation, it provides valuable insight into potential VS within individual plants (Hochberg et al. 2017; Rodriguez-Dominguez et al. 2018; Song et al. 2022). Studies using the optical method found a general trend toward lower embolism resistance in leaves than in stems, though this pattern varied among the species investigated (Cardoso et al. 2022). Because of the difficulty to measure xylem embolism resistance of below-ground organs, roots remain comparatively understudied, with only few studies addressing them (e.g. Harrison Day et al. 2024).

In this study, we used the optical method to simultaneously monitor xylem embolism formation in leaves, stems, and coarse and fine roots in eight species representing different xylem functional types and varying degrees of isohydricity. This allowed us to assess vulnerability segmentation among organs within species and compare organ-specific xylem vulnerability across species. We estimated the water potential at 12%, 50% and 88% area cavitated (P_12_, P_50_, and P_88_, respectively) in leaves and fine roots to get a measure of the classical hydraulic VS. In addition, we introduce a novel metric ‘temporal xylem vulnerability’ to quantify the elapsed dehydration time to reach 12%, 50% and 88% embolism, providing information on temporal VS. This approach accounts for environmental variability and enables comparison across species. We expressed the standardized timings as T_12_, T_50_, and T_88_ (in kPa hours until 12%, 50% and 88% area embolized). Finally, we measured minimum conductance across all four organs and eight species. We hypothesized that (1) embolism resistance decreases in the sequence stem > coarse roots > leaves/needles > fine roots; (2) the isohydric species exhibit greater degrees of segmentation than the anisohydric species; (3) the elapsed time to critical xylem embolism thresholds is positively related to more negative xylem pressures at these thresholds; and (4) the elapsed time is negatively related to the species’ minimum conductance.

## Materials and Methods

### Plant Material

The study was conducted in the outdoor, greenhouse and climate chamber facilities. We used 1-3-year-old plants sourced from the forest nursery Hermann von Appen in Soderstorf, Northern Germany. Eight temperate tree species encompassing different xylem-structural types were studied, including four conifers – *Abies alba* (European silver fir), *Picea abies* (Norway spruce), *Pinus sylvestris* (Scots pine) and *Pseudotsuga menziesii* (Douglas fir), and four deciduous broadleaved species – *Fagus sylvatica* (European beech), *Acer platanoides* (Norway maple), *Quercus robur* (Pedunculate oak) and *Tilia cordata* (Small-leaved lime). *A. platanoides* and *T. cordata* are diffuse-porous species, *Q. robur* is a ring-porous species, and *F. sylvatica* is a diffuse-porous to semi-ring-porous species (Schoch et al. 2004). All conifers studied here are generally considered isohydric (Martínez-Vilalta et al. 2014; Magh et al. 2020; Paligi et al. 2024; Schumann et al. 2024), as is maple (Leuschner et al. 2024; Kunz et al. 2016; Zeppan et al. 2026). In contrast, *F. sylvatica* and *Q. robur* are typically anisohydric, while *T. cordata* has shown to shift between iso- and anisohydry (Leuschner 2020; Leuschner and Meinzer 2024; Paligi et al. 2024; Waite et al. 2024) (see also Table 1). For each species, 15 individuals were potted at the end of March 2024 into 5 L-pots filled with sand. Plants were then placed outdoors and regularly watered (see Table 1 for species age, morphological traits, stomatal closure points and hydroscape area, used as an indicator of species isohydricity).

**Table 1.**
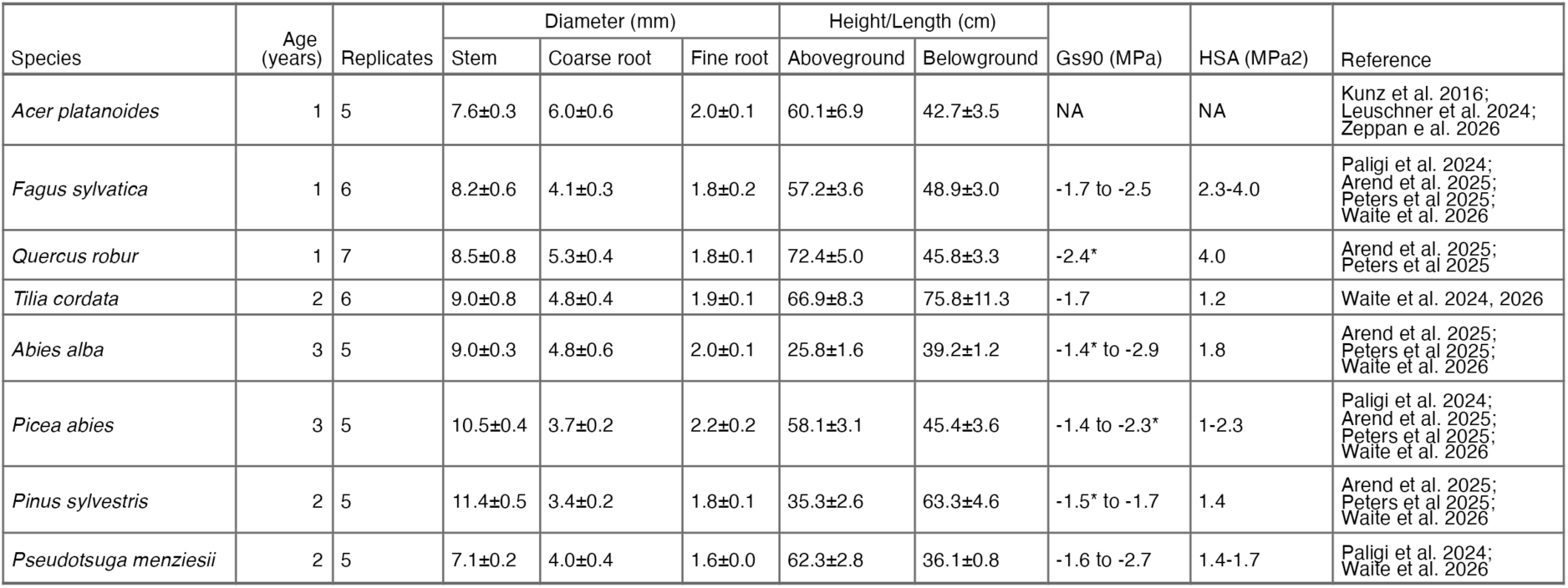
Summary of the studied tree species, their morphological traits, and hydraulic parameters characterizing species isohydricity. Morphological traits include stem, coarse-root, and fine-root diameters (mm), as well as aboveground and belowground heights (cm). Values are presented as mean ± standard error. Gs90 represents water potential at 90% stomatal closure (MPa); an asterisk (*) indicates values reported in the literature as complete stomatal closure. HSA represents hydroscape area (MPa²). Hydraulic parameters were compiled from the cited literature.

### Measurement conditions

The measurements of xylem embolism resistance were conducted between June and October 2024 using the optical method with at least five replicate plants per species. A day before the measurement, the plants were removed from the pot and the roots carefully washed, minimizing any damage to the belowground part of the plant while removing soil to facilitate installation of Cavicams on the roots. If there was any cut-open end or damage to the roots, they were glued to reduce open cavities, which would otherwise induce air entry resulting in embolism spread. After careful washing of the roots, they were soaked in water in the dark at 8 °C overnight to ensure the measurements started on a well-hydrated plant. During the day of measurements, the plants were moved to a climate chamber (BBC, Brown Boveri-York, Hamburg, Germany). Temperature (T) and relative humidity (RH) within the chamber were recorded using a HOBO MX1101 Temperature/Relative Humidity Data Logger (Onset Computer Cooperation, Bourne, MA, USA). The chamber was programmed to simulate a diurnal cycle with peak daytime conditions of 27 °C and 45% relative humidity and night-time low conditions of 15 °C and 70% relative humidity. This was done to simulate the diurnal cycle of temperature and relative humidity and to also avoid higher dehydration rates during night-time, since this would have required water potential measurements during night-time. The T and RH data was used to calculate vapor pressure deficit (VPD) every five minutes; the resulting diurnal pattern of VPD fluctuation inside the chamber is shown in Figure S1.

The resulting VPD values were multiplied with the time spans (in h) to a physiological event (elapsed time) and accumulated to obtain elapsed dehydration time weighted by the corresponding vapor pressure deficit (unit: kPa hours). Missing values from days when the data logger malfunctioned were replaced with the mean VPD recorded at the corresponding time of day from another measurement date, as similar climate chamber conditions were maintained throughout the experiment (Fig. S1).

### Optical vulnerability curves

Optical vulnerability (OV) was simultaneously measured with Cavicams (cavicam.co, Hobart, TAS, Australia) in four plant organs: fine roots, coarse roots, stems and leaves/twigs (terminal twigs in case of conifers) of the same individual (Fig. 1). Fine roots were defined as distal roots with maximum diameters usually <2.0 mm, while the adjacent thicker roots >2 mm in diameter were classified as coarse roots (range: 3.0 – 6 mm; see Table 1). In case of fine roots, OV measurements usually took place on segments of 3^rd^- or 4^th^-order roots in a hierarchical branching system, counted from the root tip.

**Figure 1.**
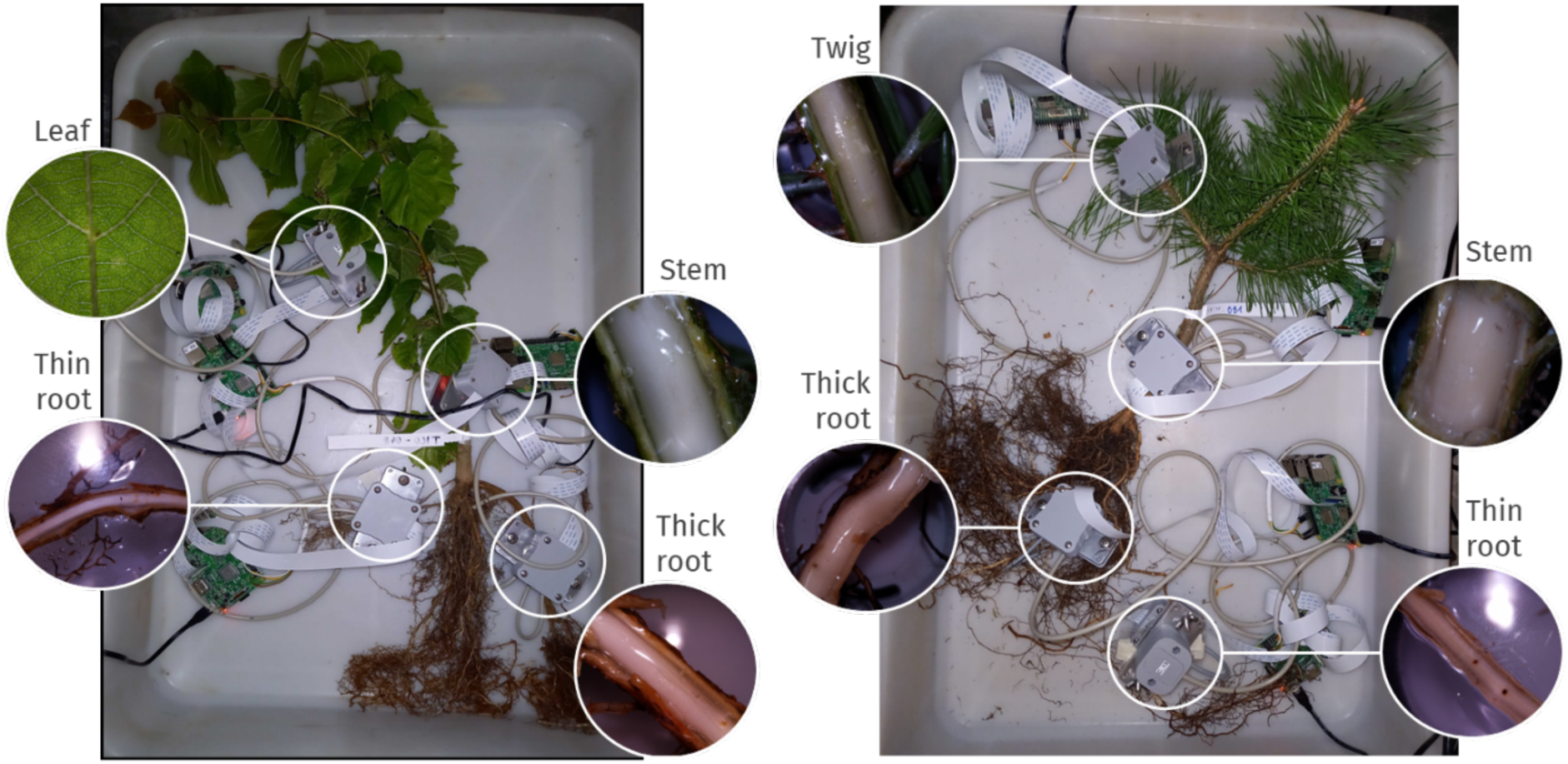
Experimental setup of Cavicams attached to different plant organs, exemplified for little-leaved lime (*Tilia cordata*; left) and Scots pine (*Pinus sylvestris;* right). Cavicams were installed on fine roots, coarse roots, leaves (broadleafs)/twigs (conifers) and stems to monitor organ-specific embolism vulnerability. Insets show close-up views of the individual camera placements on each organ.

For leaf OV measurements, one single leaf was clamped between the top and bottom plates of the imaging chamber, and transmitted light setting was used. Because conifer needles were too thick to capture embolisms with the transmitted light setting, a distal current-year twig was used instead. The Cavicam was installed on the twig after carefully removing needles and exposing the xylem with a razor blade, while ensuring that the xylem was not damaged. The exposed xylem was then covered with hydrogel (Tensive Adhesive Gel, Parker Laboratories Inc., Fairfield, NJ, USA) to prevent surface evaporation and improve light transmission (Harrison Day et al. 2024; 2026). After the twig was clamped between the two plates of the imaging chamber, the reflected light setting was used. The roots and stems were prepared similarly, first exposing the xylem, then covering the exposed xylem with hydrogel, and using the reflected light setting. Additionally, the fine root was fixed between two glass slides to prevent distortion due to drying of the root cortex. Before removing the bark, phloem and cambium of the installation sites of the roots and stems, the diameter of the segments was measured using a digital caliper (Horex. (n.d.). Digital caliper 200 mm). The shutter speed was adjusted for each installation before starting measurements. The plants dried naturally on a bench, while images were captured at five-minute-intervals, and water potential measurements on leaves/twigs and fine roots were taken in parallel using a Scholander pressure chamber.

### Water potential measurements

The xylem water potential of fine roots and foliage was measured at intervals of 1-8 h, depending on the degree of hydration, from full saturation to complete dehydration using a Scholander pressure chamber (PMS instruments, Corvalis, OR, USA). For leaf water potential measurements, a single leaf/needle (i.e., in *A. platanoides*, *T. cordata*, and *P. sylvestris*) or terminal twig was used. For root water potential measurements, fine roots ∼2 mm in diameter were chosen to match the size of those used for Cavicam installation. The cut leaf/fine root area was sealed with glue to prevent excessive dehydration and subsequent embolism propagation. Before measurement, the sample was wrapped in aluminium foil and was allowed to equilibrate for 15 to 60 min (Rodriguez-Dominguez et al. 2022). Because leaf and fine root water potentials were measured periodically, we applied a shape-constrained additive model using the R package `SCAM v1.2.20` (Pya 2025) to interpolate water potential at 5-min intervals, matching the timing of Cavicam image acquisition. A penalized monotonic decreasing spline with a basis dimension of five was fitted for each measured individual at organ level, with measured water potentials as a function of elapsed dehydration time (kPa hours; Fig. S2).

### Image processing and calculations

Image sequences were processed in ImageJ using the OSOV Toolbox (https://www.opensourceov.org/). The analysis is based on the principle that gas-filled vessels reflect more light than water-filled vessels (Haines 1935), resulting in detectable pixel-changes between two sequential images when an embolism forms.

Residual noise and artefacts caused by tissue shrinkage, movement of the plant material, or insects moving were removed. After this cleaning step, the remaining pixel changes were assumed to represent embolism events, which were then quantified (Fig. S3). The processed output provided the cavitated area in pixels for each image slice. Due to incomplete image sequences, some measurements were excluded from the analysis, resulting in 77 water potential-based vulnerability curves (Fig. S4) and 162 time based-vulnerability curves (Fig. S5).

### Minimum conductance

Minimum epidermal conductance (Gmin) was measured between mid and late September 2024, with four replicates per species and organ. Sample collection, transport, and storage followed the same procedures as described for the Cavicam measurements. After saturation, single leaves (in case of broadleaved species), current-year shoots (in case of conifers), and 5-cm segments of stems, coarse roots, and fine roots were excised. The open cut surfaces of each sample were sealed with high-melting-point wax to prevent water loss through the cut ends of the xylem. Samples were placed in a climate chamber with limited light exposure (BBC, Brown Boveri-York, Hamburg, Germany) maintained at 25 ± 0.5 °C, 50 ± 2% relative humidity. Sample mass was repeatedly measured using an analytical balance (0.1 mg precision) until mass loss approached a constant rate (Kerstiens 1996; Sack et al. 2003). Projected leaf and needle area were determined before the measurement using WinFOLIA, while stem and root surface areas were quantified using WinSEEDLE (Régent Instruments Inc., Quebec, Canada). Following the measurements, samples were oven-dried at 70 °C for approximately 72 h to determine dry mass. Relative water content (RWC) was calculated throughout dehydration using sample fresh mass, dry mass, and saturated mass.

For intervals exceeding 20 min between consecutive measurements, sample mass was interpolated as a function of dehydration time using shape-constrained additive models (SCAM; Pya 2025). Gmin was estimated from the slope of the linear relationship between water loss and time within four RWC levels (95–85%, 85–75%, 75–65%, and 65–55% RWC). Conductance estimates were subsequently standardized by sample surface area and expressed in mmol m^−2^ s^−1^.

### Statistical analysis

Data and statistical analysis were carried out in `R v4.2.0 (`R Core Team 2022) under the framework of the `tidyverse v2.0.0` R package (Wickham et al. 2019).

The cumulative cavitated area was calculated from image sequences and scaled to obtain the percentage cavitated area over the course of dehydration. Relationships between percentage embolism and both xylem water potential (Ψ; for leaves/twigs and fine roots) and elapsed dehydration time (kPa hours; for leaves/twigs, fine roots, coarse roots and stem) were modelled separately for each species and organ combination for each individual Cavicam installation using shape-constrained additive models in R package `scam v 1.2-20` (Pya 2025). Models were fitted with a monotonic increasing P-spline smoother (bs = “mpi”) and a Gamma error distribution with log link function. A smoothing parameter (sp = 0.001) and gamma penalty of 1.4 were applied (Fig S4, S5). From the fitted curves, we applied the linear interpolation method using the `approx` function in R to extract the xylem water potentials corresponding to 12%, 50%, and 88% cavitated area (denoted as P_12_, P_50_, and P_88_, respectively) and elapsed dehydration time corresponding to 12%, 50%, and 88% cavitated area (denoted as T_12_, T_50_, and T_88_, respectively).

Differences in modelled thresholds among species, organs, and embolism parameters were tested using generalized linear mixed-effects models (glmer, `lme4` package) with Gamma error distribution and log link (Bates et al. 2014). Fixed effects included plant organ (testing hypothesis 1: distal organs are more sensitive than proximal organs), embolism threshold (12, 50, 88% embolized areas), species (testing hypothesis 2: species with higher degree of isohydricity show greater segmentation), and their interactions, while plant identity was included as a random effect with random slopes for organ and parameter. A Spearman rank correlation test to test was conducted to test hypothesis 3: whether more negative water potentials are associated with longer dehydration times to 12%, 50% and 88% embolism (P_12_, P_50_, and P_88_, vs. T_12_, T_50_, and T_88_).

Similarly, GLMMs were employed to test for differences in minimum conductance (G_min_) among organs (fine root, coarse root, leaf/twig and stem), dehydration intervals (RWC95–85, RWC85–75, RWC75–65, and RWC65–55), species, and their interactions. Plant identity was included as a random effect with random slopes for organ and parameter, and models were fitted with a Gamma error distribution and log-link function. For hypothesis 4, we tested whether minimum epidermal conductance predict time to embolism thresholds using a linear mixed-effects model with time to embolism threshold as the response and minimum epidermal conductance, organ, and embolism threshold as interacting predictors, including relative water content category as a covariate and species as a random intercept. The model performance was evaluated using the `check_model` and `compare_performance` functions from the `performance v0.13.0` R package (Lüdecke et al. 2021). Post-hoc pairwise comparisons for all models were conducted with the `emmeans` function from the `emmeans v1.9.4.1` R package (Lenth et al. 2023) to obtain comparisons between species and among species and different organs of the plant. All figures were produced using the `ggplot v3.5.1` package (Wickham 2016), and tables using the `flextable v0.9.7` R package (Gohel and Skintzos 2017).

## Results

### Critical xylem water potential thresholds

#### Leaf xylem vulnerability thresholds across species

Among leaves/twigs, *Abies alba* followed by *Quercus robur* consistently exhibited the greatest embolism resistance, with the most negative P_12_, P_50_, and P_88_ values (Table S2). Leaf P_50_ in *A. alba (*- 4.69 MPa) was significantly more negative than in *Picea abies* (-2.05) and *Pinus sylvestris* (-2.14), while leaf P_88_ was significantly more negative than in *P. abies* (P < 0.05; Table S2; Fig. S6). In contrast, *P. abies* and *T. cordata* generally showed comparatively vulnerable leaves with less negative thresholds, whereas *Q. robur, Fagus sylvatica* and *Pseudotsuga. menziesii* revealed intermediate values that were comparable across thresholds (Fig. S7).

#### Fine-root xylem vulnerability thresholds across species

Species differences were equally pronounced in fine roots as in leaves. Fine-root P_50_ values ranged from -0.97 MPa in *P. sylvestris* to -4.8 MPa in *A. alba*. *P. abies, T. cordata* and *P. sylvestris* consistently showed vulnerable fine roots, characterized by the highest (least negative) P_12_, P_50_, and P_88_ values, whereas *A. alba* followed by *Q. robur* and *A. platanoides* showed the greatest resistance (Table S2). Vulnerability thresholds in *A. alba* were significantly more negative than *P. abies, T. cordata, P. sylvestris and F. sylvatica*, while *Q. robur* and *A. platanoides* also showed significantly more negative fine-root P_12_ than these four species (P < 0.05; Table S2, Fig 2).

**Figure 2.**
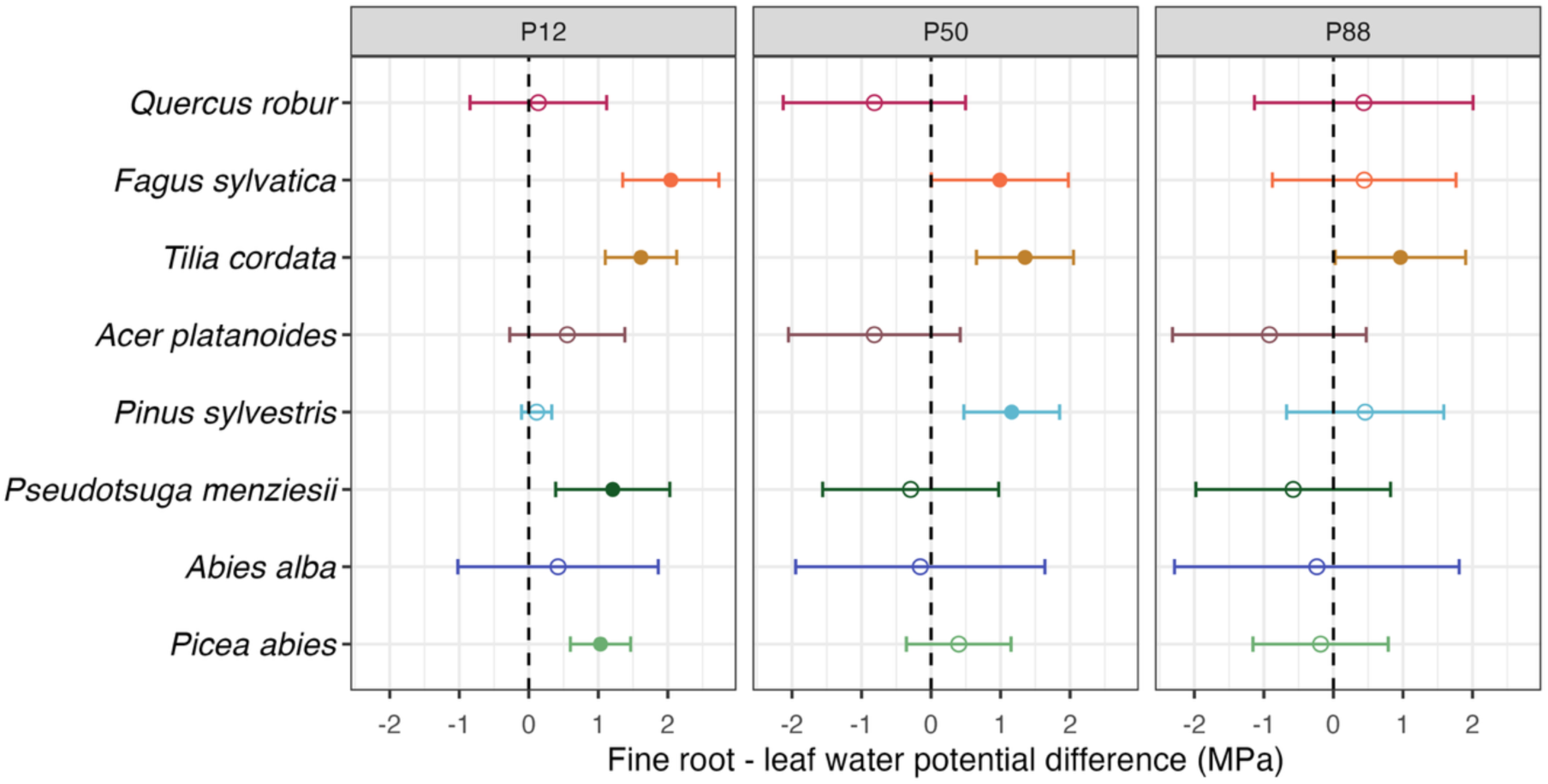
Difference in the P_12_, P_50_ and P_88_ values (water potentials of 12, 50 or 88 % loss of hydraulic conductance) of fine root and leaf/needle xylem in the eight species, given as estimated marginal means of the xylem water potential difference. Species are ordered from least to more isohydric top to bottom in y-axis. Points represent the estimated marginal mean differences between fine roots and leaves/needles, with 95% confidence intervals indicated by the bars. Positive differences indicate less negative xylem water potentials in fine roots than in leaves. Closed circle indicates significant differences from zero and open circles non-significant differences. The number of replicate measurements varied among species-organ combinations (*n* ≥ 3, typically *n* > 4 & < 7).

### Elapsed time to reach water potential thresholds across species

Elapsed dehydration time to critical embolism thresholds (T_12_, T_50_ and T_88_) differed strongly among species and plant organs (Table S3, S4). Species differences were more pronounced in stems and became increasingly pronounced toward higher embolism thresholds. At T_12_, *P. sylvestris* consistently exhibited shorter stem dehydration times than most other species, differing significantly from *A. platanoides, F. sylvatica, Q. robur,* and *T. cordata* by ∼12-23 kPa hours (Fig. S8; Table S4). Similar patterns persisted at T_50_, where stems of *A. alba, A. platanoides*, and *F. sylvatica* maintained dehydration times 25-38 kPa hours longer than *P. sylvestris*. At T_88_, the strongest species contrasts occurred in stems. *A. alba* and *F. sylvatica* maintained the longest dehydration times, while *P. menziesii* and *P. sylvestris* reached embolism thresholds substantially earlier. Stem dehydration times in *A. alba* (by ∼37-40 kPa hours) and *F. sylvatica* (by ∼26-29 kPa hours) exceeded those of *P. sylvestris* and *P. menziesii* (P < 0.001; Fig. S9; Table S4).

Species differences in leaves and fine roots were generally weaker but still evident at lower embolism thresholds. In leaves at T_12_, *Q. robur* showed significantly longer dehydration times than *P. abies* and *P. sylvestris* (∼13 kPa hours; P < 0.05), while *A. platanoides, A. alba* and *T. cordata* differed at marginal significance (P < 0.1). In fine roots at T_12_, *Q. robur* showed consistently longer dehydration times than *P. sylvestris, T. cordata,* and particularly *F. sylvatica* (by ∼13-22 kPa hours; P < 0.01; Table S4), while for T_50_, only *P. sylvestris* differed with marginal significance from *Q. robur*. In contrast, coarse roots showed no statistically significant differences among species, despite considerable variation in the timing of hydraulic failure (Table S4, Fig S7). For example, *Q. robur* reached its temporal threshold markedly later (∼13-36 kpa hours) than the other studied species.

### Xylem vulnerability segmentation between fine roots and leaves

Hydraulic segmentation between fine roots and leaves varied substantially among species, and was different for P_12_, P_50_ and P_88_. Overall, organ differences were greatest at P_12_ and P_50_, and decreased toward P_88_ (Table S1; Fig. 2). Four of the eight studied species exhibited significant organ-level differences in vulnerability thresholds, particularly at P_12_, and P_50_ (Table S1). In *P. sylvestris*, *T. cordata*, *F. sylvatica* and *P. abies,* leaves tended to be more embolism-resistant than fine roots, although the magnitude of differences varied across thresholds. In contrast, organ differences were generally smaller and mostly non-significant in *A. alba, A. platanoides*, *P. menziesii* and *Q. robur*, with fine roots tending to show more embolism resistance compared to leaves at P_50_ and P_88_ (Fig. 2, S10; Table S1).

### Time line of vulnerability segmentation across plant organs

Elapsed dehydration time to reach a critical embolism threshold differed markedly among the plant organs of a species and differences between organs became more pronounced at T_50_ and T_88_ than at T_12_, indicating an increasingly different timing of the breaching of embolism thresholds in the organs with progressive dehydration (Fig. 3, Fig S8, S9; Table S3). Overall, stems tended to take longer time to reach a critical embolism threshold than other organs (few exceptions occurred in *P. menziesii*, *P. sylvestris* and *Q. robur*), with fine roots typically reaching critical embolism thresholds fastest (Table S3; Fig. 3, S11). *F. sylvatica* showed the strongest vulnerability segmentation. At T_88_, fine roots, leaves and coarse roots reached embolism thresholds ∼27-30 kPa hours earlier than stems (P < 0.01), while similarly strong differences were evident at T_50_ by reaching critical embolism thresholds 18-24 kPa hours earlier (P < 0.05). *A. alba* showed a comparable pattern, particularly at T_88_, where fine roots and coarse roots dehydrated 26-29 kPa hours faster than stems (P < 0.05), while marginally significant differences were also observed at T_50_ for both organs. *A. platanoides* stems reached embolism thresholds ∼15 kPa hours later at T_50_ and T_88_ than leaves (P < 0.05), while fine roots differed only at T_12_. Similarly, *T. cordata* also showed segmentation, with fine-roots and coarse roots reaching T_12_ and T_50_ ∼10-15 kPa hours earlier than the stem (P < 0.05).

**Figure 3.**
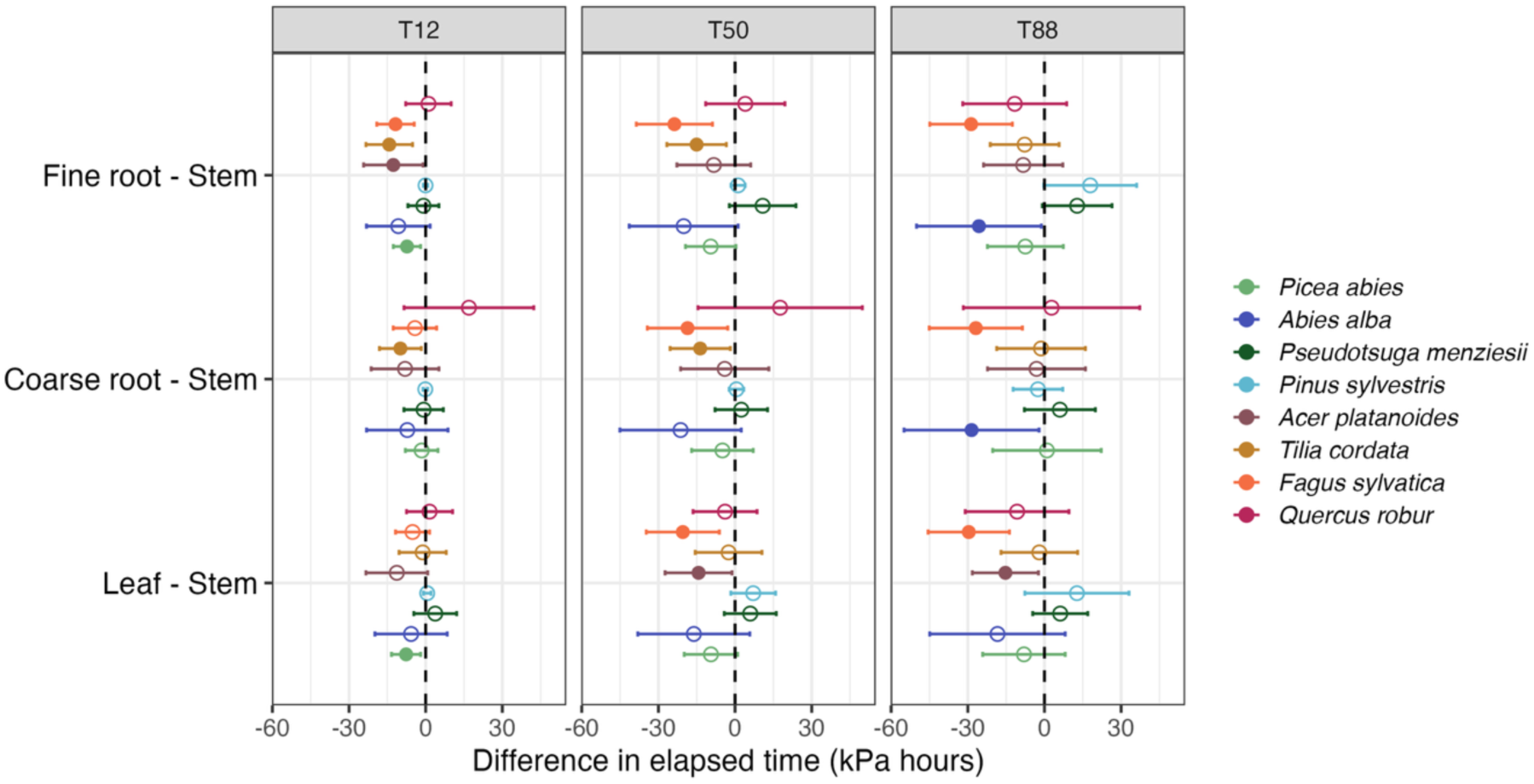
Difference in the T_12_, T_50_ and T_88_ values (time in kPa hours elapsed to reach water potentials of 12, 50 or 88 % loss of hydraulic conductance) between fine root and stem xylem (upper block), between leaf/needle and stem xylem (central block), and between coarse root and stem xylem (lower block) in the eight species, given as estimated marginal means of the length of the time interval. Species are ordered from least to more isohydric top to bottom in y-axis. Points represent the estimated marginal mean differences between the each two organs, with 95% confidence intervals indicated by the bars. Positive differences indicate longer dehydration times in fine roots, leaves or coarse roots than in stems to reach the P_12_, P_50_ or P_88_ thresholds. Closed circle indicates significant differences from zero and open circles non-significant differences. The number of replicate measurements varied among species-organ combinations (*n* ≥ 3, typically *n* > 4 & < 7).

In contrast, *P. abies* displayed weak vulnerability segmentation, with significant differences only at T_12_, where fine roots and leaves dehydrated ∼8 kPa hours earlier than stems. *P. sylvestris* and *P. menziesii* showed the opposite tendency at T_88_, where fine roots tended to maintain longer dehydration times than stems (difference marginally significant at P < 0.1; Fig. 3; Table S3). *Q. robur* showed the weakest segmentation among the eight species with coarse roots generally taking longer to reach embolism thresholds than the other organs. Temporal differences between fine and coarse roots varied little among species and thresholds (Fig. 4).

**Figure 4.**
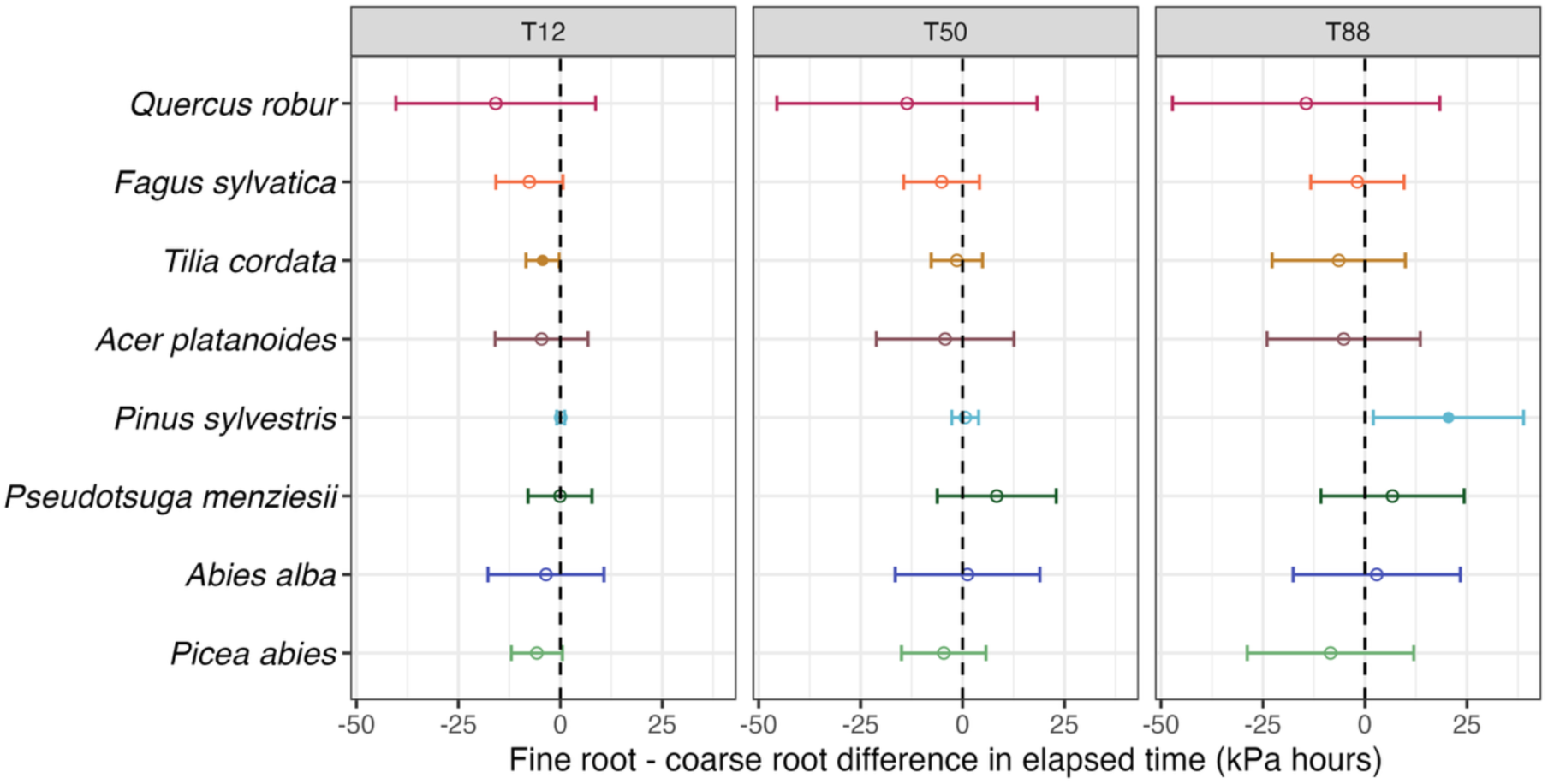
Difference in the T_12_, T_50_ and T_88_ values (time in kPa hours elapsed to reach water potentials of 12, 50 or 88 % loss of hydraulic conductance) between fine root and coarse root xylem in the eight species, given as estimated marginal means of the length of the time interval. Points represent the estimated marginal mean differences between the fine root and coarse root xylem, with 95% confidence intervals indicated by the bars. Positive differences indicate longer dehydration times in fine roots than coarse roots to reach the P_12_, P_50_ or P_88_ thresholds. Closed circle indicates significant differences from zero and open circles non-significant differences. (*n* ≥ 3, typically *n* > 4 & ≤ 7).

### Correlation between xylem water potential and elapsed dehydration time causing embolism

Correlations between xylem water potential and elapsed dehydration time varied considerably among species, organs, and embolism thresholds (Fig. 5). Overall, more negative water potentials were generally associated with longer dehydration times, resulting in predominantly positive correlations in both fine roots and leaves (Fig. 5, Fig S12). Across species, correlations were generally more positive in leaves than in fine roots.

**Figure 5.**
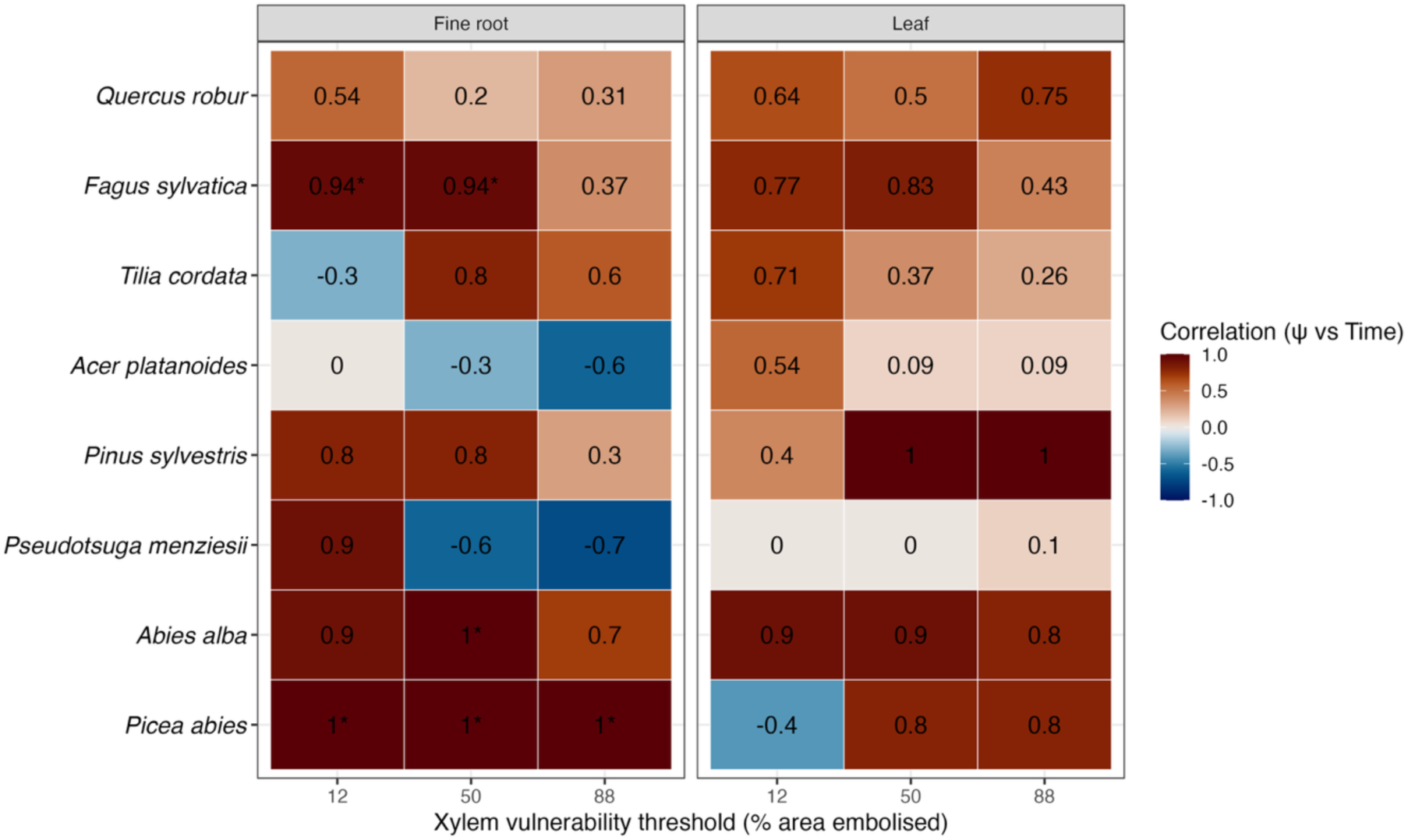
Heatmap graph showing the Spearman’s rank order correlation coefficients for the relations between the P_12_, P_50_ and P_88_ values (water potential thresholds) and the corresponding T_12_, T_50_ and T_88_ values (time spans needed to reach P_12_, P_50_ and P_88_) in the eight species, given separately for fine roots (left block) and leaves/needles (right block). Correlations were calculated separately for fine roots and leaves. Color intensity indicates the strength and direction of the relationship, with blue representing negative correlations and red representing positive correlations. Positive correlations indicate that more negative xylem water potentials are associated with longer elapsed dehydration time, whereas negative correlations indicate the opposite trend. Asterisks denote statistically significant correlations (P < 0.05). (*n* ≥ 3, typically *n* > 4 & ≤ 7). Non-significant results should be interpreted cautiously given the limited sample size, despite perfect correlations (rₛ = 1).

The strongest and most consistent positive correlations occurred in *P. abies, F. sylvatica,* and *A. alba*, particularly in fine roots (with correlation coefficients > 0.9 across all thresholds). Leaves of *P. sylvestris* and *A. alba* also showed strong positive relationships, especially at higher embolism thresholds (P_50_ and P_88_). In contrast, *P. menziesii* and *A. platanoides* displayed weaker or negative correlations in fine roots at higher embolism thresholds, where *T. cordata* showed contrasting responses between thresholds, with weaker relationships at P_12_ than at P_50_ and P_88_.

### Organ-specific variation in minimum conductance

Dehydration patterns were strongly organ-dependent in all species. Stems consistently displayed the slowest RWC decline, whereas fine roots dehydrated most rapidly followed by coarse roots. Leaves and needles exhibited intermediate dehydration rates (Fig. 6, S13, S14). Minimum conductance (Gmin) varied systematically among organs and dehydration stages. Across all RWC levels examined, fine and coarse roots generally exhibited higher Gmin than stems (differences significant in all species and dehydration levels; Fig. 6, S15; Table S5). The largest root-stem contrasts occurred in the deciduous species *F. sylvatica, A. platanoides*, and *T. cordata*, whereas conifers showed smaller organ-level differences and a generally lower root Gmin (Fig. S13, S14).

**Figure 6.**
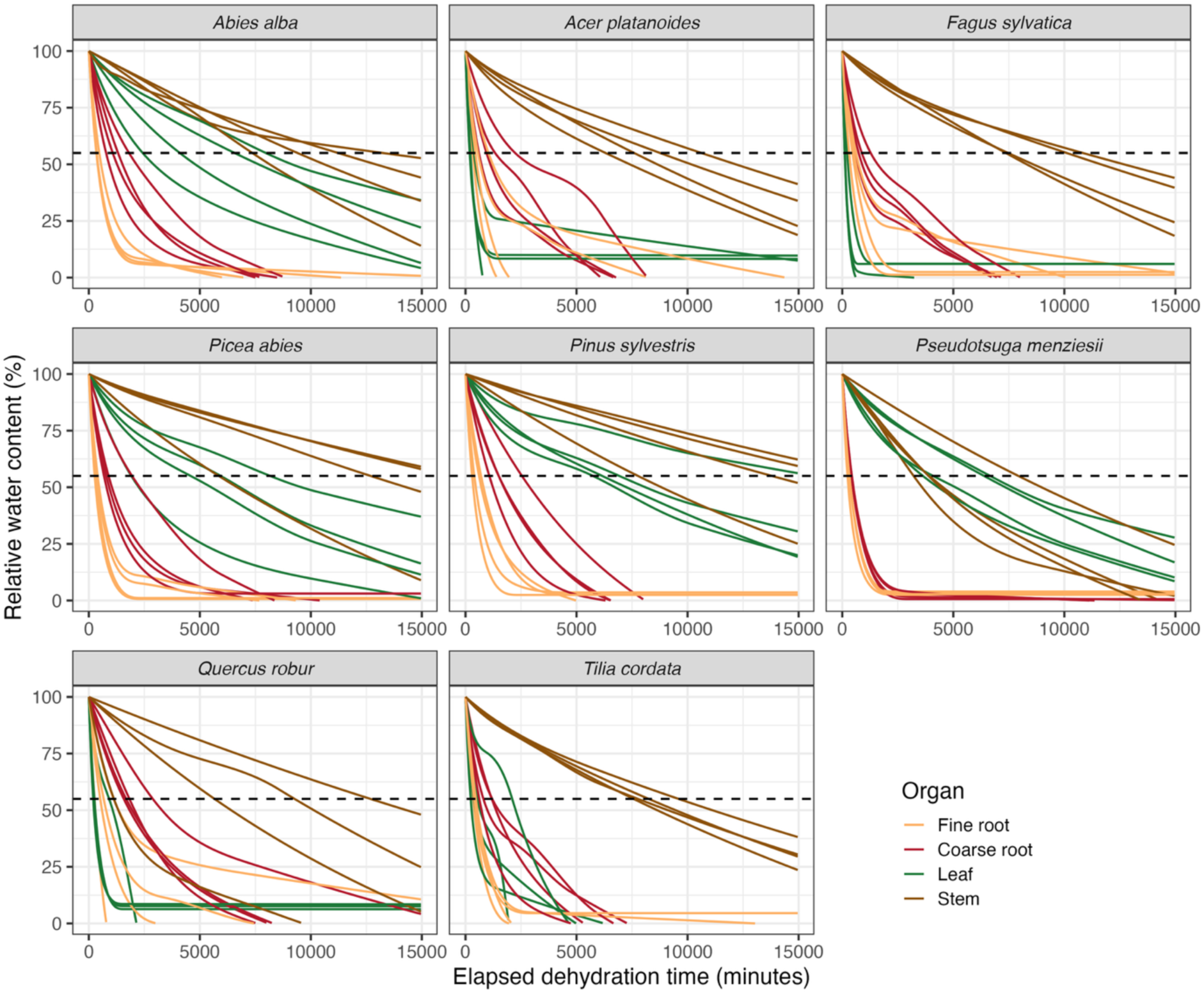
Relative water content (RWC, %) decline during dehydration of four plant organs (fine roots, coarse roots, leaves/needles, and stems) in the eight studied tree species. Curves represent individual samples monitored throughout dehydration, and colors distinguish organ types (orange = fine roots, red = coarse roots, green = leaves/needles, brown = stems). The horizontal dashed line indicates 55% relative water content, shown as a reference threshold. The x-axis represents elapsed dehydration time (minutes) and the y-axis relative water content (%). Sample sizes varied among species and organ combinations (generally n ≥ 3 & n ≤ 4).

In correspondence, conifers and broadleaved species differed in the leaf-stem contrasts in Gmin. In all four conifer species, needles consistently exhibited significantly lower Gmin than stems across all RWC levels (P < 0.05; Fig. S16; Table S5). In contrast, leaf and stem Gmin did not differ significantly in the four broadleaved species. Across species, Gmin declined with decreasing RWC, but the relative ranking of organs remained largely unchanged, with roots generally exceeding stems and leaves (Fig. 6, S15).

### Relationship between minimum conductance and embolism resistance

Across organs and tree functional groups, higher Gmin was generally associated with shorter dehydration times to reach embolism thresholds (Fig. 7). However, the strength of this relationship varied among organs, functional groups, and embolism thresholds.

**Figure 7.**
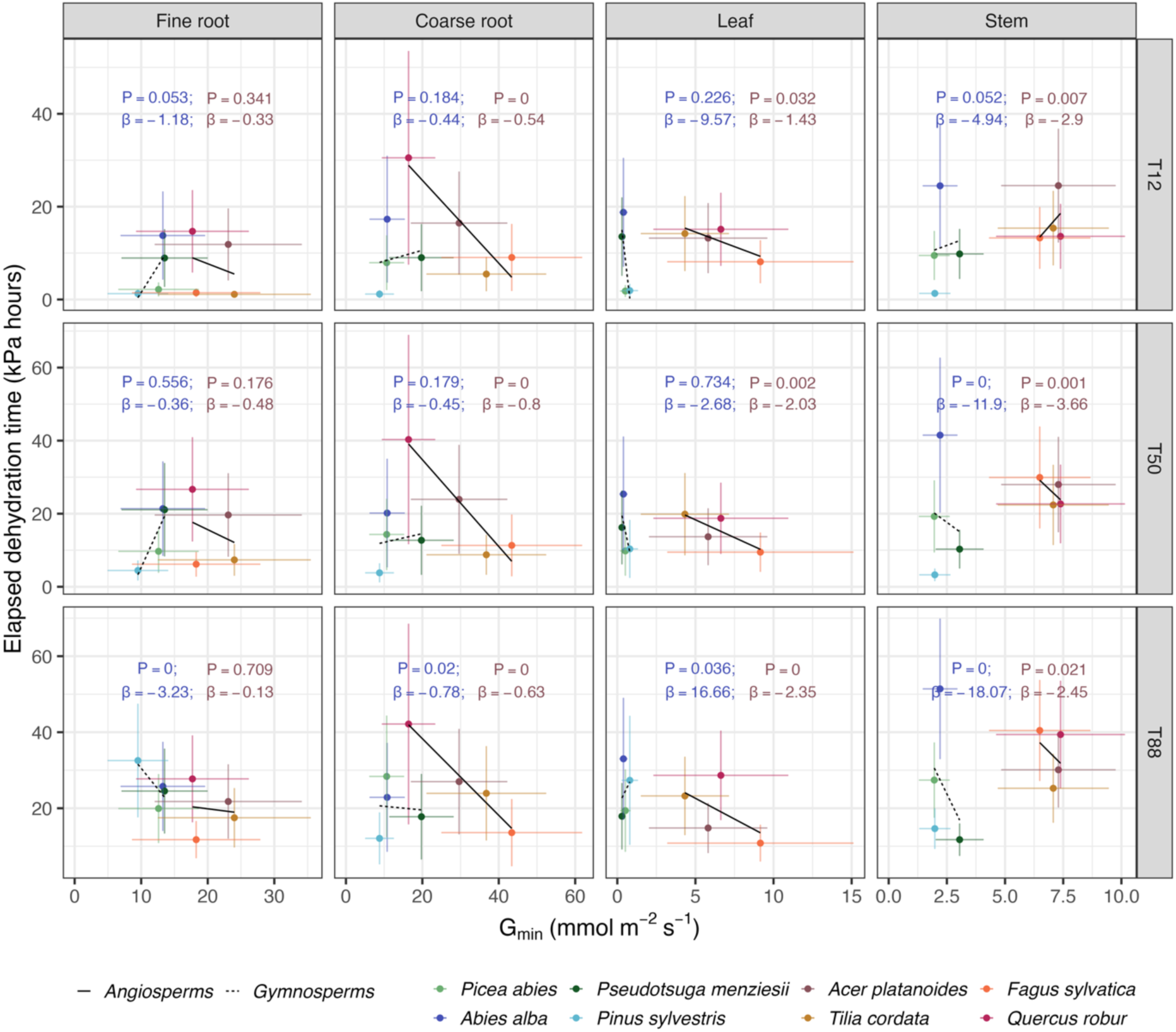
Relationships between minimum conductance (Gmin) and elapsed dehydration time to embolism thresholds (T_12_, T_50_, and T_88_) for fine roots, coarse roots, leaves, and stems in the eight species. Differences between gymnosperms and angiosperms are highlighted by dashed vs solid lines. Points represent species means and their 95% CI of Gmin or elapsed time to embolism threshold. Separate least-squares regression lines were fitted for angiosperms (solid black lines) and gymnosperms (dashed black lines) within each organ and embolism threshold. Reported regression statistics are from a linear mixed effect model. Within each panel, regression slope (β), and p-value for each functional group (blue, gymnosperms; brown, angiosperms) are shown. Positive slopes indicate longer dehydration times with increasing Gmin, whereas negative slopes indicate shorter dehydration times.

In the broadleaved species, Gmin was consistently negatively associated with dehydration time in leaves, coarse roots, and stems. The slope between dehydration time and Gmin ranged for leaves from -1.43 to -2.35 kPa h per mmol m^-2^ s^-1^ across T_12_–T_88_ (all P ≤ 0.032). Coarse roots exhibited weaker but highly consistent negative slopes (-0.54 to -0.80; all P < 0.001), whereas stems showed the steepest declines in dehydration time with increasing Gmin among the organs of the broadleaf species (−2.45 to −3.66; P < 0.03). In contrast, fine root dehydration time showed no significant relationship with Gmin (P > 0.05; Fig. 7; Table S6).

In the conifers, responses were more organ- and threshold-dependent. Stem Gmin exhibited the strongest negative Gmin – dehydration time relationships, with slopes becoming progressively steeper from T_12_ (-4.94; P = 0.052) to T_50_ (-11.90; P < 0.001) and T_88_ (-18.07, P < 0.001). Fine roots were negatively associated with dehydration time only at T_88_ (-3.23, P < 0.001), with a marginal effect at T_12_ (-1.18, P = 0.053), while coarse roots showed a weaker negative relationship only at T_88_ (-0.78, P = 0.020). In contrast, needles showed no significant relationship at T_12_ or T_50_ (P > 0.22), but exhibited an unexpected positive association at T_88_ (16.66, P = 0.036).

## Discussion

In this study, we quantified hydraulic and temporal dimensions of xylem vulnerability across leaves, stems, coarse roots, and fine roots in eight temperate tree species. Both embolism resistance and the time required to reach critical embolism thresholds varied substantially among organs and species and were not consistently related to the species’ isohydricity. Embolism resistance (P_12_, P_50_ and P_88_) and vulnerability dynamics (T_12_, T_50_, T_88_) were often decoupled. While some species exhibited pronounced vulnerability segmentation among organs, others showed more coordinated hydraulic behaviour. Furthermore, minimum conductance (Gmin) emerged as an important predictor of dehydration timing in the broadleaf trees, whereas no consistent relationship was observed in the conifers.

### Organ-level vulnerability segmentation is not related to species isohydricity

Xylem vulnerability was highly organ- and species-specific and did not conform to a consistent pattern across the eight species studied. In terms of water potential-based thresholds, leaves tended to exhibit more negative thresholds than fine roots in several species, particularly *F. sylvatica* and *T. cordata*. In contrast, other species showed little differentiation between organs in vulnerability levels, and *A. platanoides* and *Q. robur* exhibited for P_50_ an opposite tendency, i.e. fine roots possessing by ∼0.7 MPa more negative P_50_ values than leaves (though the differences were non-significant; Fig. 2 and Table S1). This variability suggests that hydraulic vulnerability segmentation is not governed by a universal hydraulic vulnerability hierarchy between leaves and roots, but may instead reflect species-specific hydraulic strategies. For instance, greater leaf resistance may allow leaves to maintain hydraulic function under increasingly negative water potentials, whereas greater root resistance could help sustain water uptake, as soil water availability declines.

Beyond differences in the water potential at which embolism thresholds were reached, species also differed in the time required to reach these thresholds, revealing an additional dimension of plant vulnerability to drought. Five of the eight species exhibited clear temporal vulnerability segmentation, with stems generally requiring longer to reach critical embolism thresholds than coarse roots, leaves and fine roots, particularly at P50 and P88 (Fig. 3). This pattern may reflect differences in organ structure and anatomy, hydraulic architecture and water transport dynamics, whereby woody stems may exhibit slower progression of embolism or greater hydraulic buffering compared with more distal organs (Meinzer et al. 2003; Scholz et al. 2007; Preisler et al. 2022). Such buffering can slow declines in water potential and delay embolism progression during drought, even when xylem embolism resistance itself differs little among organs (Holbrook and Sinclair 1992; Meinzer et al. 2008). The longer time required for stems to reach critical embolism thresholds could therefore indicate a temporal separation in hydraulic deterioration among organs, potentially allowing distal organs to experience hydraulic dysfunction before more structurally and functionally persistent parts of the hydraulic pathway are compromised.

However, this pattern was not universal. In *Q. robur*, coarse roots were the most resistant requiring longer time to critical thresholds. This prolonged maintenance of root hydraulic function of oak roots aligns with the well-documented drought resilience and resprouting capacity of oaks following canopy dieback (Lloret et al. 2004; Huo et al. 2024). In *P. sylvestris*, leaves and fine roots were more resistant than stems, whereas in *P. menziesii*, fine roots, followed by leaves and coarse roots, were more resistant over time than stems at P50 and P88. These exceptions indicate that temporal vulnerability segmentation is also species-dependent and cannot be explained simply by organ position or structural permanence.

Taken together, our findings do not support our hypothesis 1 which adopts the classical vulnerability segmentation hypothesis and predicts that distal organs act as hydraulic fuses and experience hydraulic dysfunction before more permanent tissues do, thereby protecting the whole-plant hydraulic function during drought (Johnson et al. 2016; Cuneo et al. 2016; Hochberg et al. 2017). Although segmentation was observed, no consistent hierarchy of vulnerability among organs was found. A similar variability in vulnerability patterns has been detected in recent optical and microCT studies, which reported no, conventionally expected or reversed trends in xylem vulnerability (Rodriguez-Dominguez et al. 2018; Peters et al. 2020; Li et al. 2020; Wilkening et al. 2023; Harrison Day et al. 2024, Peters and Choat 2025). The absence of a consistent pattern across species suggests that different strategies of coordinating hydraulic vulnerability across the plant does exist, rather than a single optimal distribution of hydraulic risks cross organs.

Furthermore, interspecific variation in the segmentation of hydraulic vulnerability and vulnerability dynamics was not consistently related to the species’ degree of isohydricity, as deduced from the stringency of stomatal regulation (Wason et al. 2018; Losso et al. 2019; Hartmann et al. 2019; Leuschner et al. 2019; Leuschner 2020; Leuschner and Meinzer 2024). Vulnerability segmentation was observed in several species conventionally classified as isohydric, notably *A. alba, A. platanoides, P. abies* and *T. cordata*, but was absent or less pronounced in the markedly isohydric *P. menziesii* and *P. sylvestris*. Conversely, the anisohydric *F. sylvatica* exhibited significant segmentation (Figs. 2 and 3; Tables S1 and S3). Thus, our results do not support hypothesis 2, which predicted stronger segmentation in isohydric species and little or no segmentation in anisohydric species. The absence of a consistent relationship may reflect the fact that isohydry primarily characterizes a species’ water-use strategy, particularly the regulation of stomatal conductance and plant water potential under declining soil water availability, rather than its intrinsic resistance to hydraulic dysfunction (Johnson et al. 2018; Kahmen et al. 2022; Waite et al. 2024). This interpretation is consistent with recent studies reporting absent or weak relationships between stomatal regulation and xylem vulnerability segmentation (Lübbe et al. 2022; Li et al. 2024; Huo et al. 2024; Peters et al. 2024). Overall, our results support the emerging view that vulnerability segmentation is not a universal property of plant hydraulic systems, and, in consequence, whole-plant drought responses cannot be inferred from the species’ isohydricity or xylem vulnerability measured in a single organ alone.

### Decoupling between xylem water potential and elapsed dehydration time causing embolism

Although embolism thresholds and dehydration time were generally correlated, the strength and direction of these relationships varied among species and organs (Fig. 5, Fig. S12). Organs with similar water potential-based embolism thresholds often differed substantially in the time required to reach critical thresholds, and temporal differentiation was evident even where hydraulic resistance was comparable (Figs. 2 and 3; Tables S1 and S3). This contradicts hypothesis 3, which predicted a positive relationship between embolism resistance and dehydration time, and suggests that intrinsic embolism resistance alone does not determine how rapidly organs progress towards hydraulic failure. These findings highlight the importance of considering not only embolism resistance but also dehydration dynamics, as traits influencing water loss and water storage may affect the rate at which tissues approach critical hydraulic thresholds (Blackman et al. 2019; Challis et al. 2022; Copie et al. 2025).

This divergence reflects the fact that water potential-based and time-related thresholds capture different aspects of hydraulic vulnerability. While water potential thresholds define the tension in the xylem at which a certain degree of embolism is reached, time-related thresholds characterize how rapidly this state develops under the given experimental conditions. Consequently, an organ may reach a given degree of embolism at a more negative water potential, but reach that threshold relatively rapidly, while another organ can exhibit a less negative potential threshold but take longer to reach it. Thus, the temporal progression of embolism provides complementary information on hydraulic dysfunction by capturing dehydration dynamics that cannot be inferred from embolism resistance alone.

In the broadleaved species, higher Gmin was generally associated across organs with shorter times to reach embolism thresholds (Fig. 7), suggesting that residual water loss after stomatal closure can accelerate tissue dehydration and the progression towards hydraulic failure, in support of hypothesis 4. Similar relationships have been reported between minimum leaf conductance, plant dehydration time, and drought survival (Paligi et al. 2024; Waite et al. 2024; Camarero et al. 2026). Because stomata typically close before substantial embolism develops (Hochberg et al. 2017; Creek et al. 2020; Sorek et al. 2022), water loss through incompletely sealed stomata and the cuticle may become increasingly important in determining dehydration rates during the later stages of tissue dehydration (Kerstiens, 1996; Duursma et al. 2019; Camarero et al. 2026; Liang et al. 2026). In contrast, relationships between Gmin and dehydration duration were weaker and less consistent in conifers (Fig. 7). This may reflect differences in the mechanisms governing water loss and hydraulic resistance between angiosperms and gymnosperms. Conifer xylem is characterized by torus–margo pit membranes, which are typically more embolism resistant per unit of water transport efficiency than the homogeneous pit membranes found in angiosperms (Bailey, 1913; Pittermann et al. 2010; Jansen et al. 2026), while their needle-shaped leaves and thick cuticles generally limit residual water loss after stomatal closure more effectively (Paligi et al. 2024; Trueba et al. 2026). Thus, the dehydration dynamics of conifers may depend more on xylem anatomical properties, which could reduce the importance of Gmin to dehydration duration.

Together, these findings suggest that the progression towards hydraulic failure is governed not only by intrinsic embolism resistance but also by the rate at which tissues lose water following stomatal closure (Liang et al. 2026; Camarero et al. 2026). Integrating hydraulic resistance with the temporal progression of dehydration may consequently provide a more comprehensive understanding of organ-level drought vulnerability and help explain why species with similar embolism thresholds can differ in drought survival and recovery.

### Vulnerability segmentation and its role in drought resistance

Whether vulnerability segmentation (VS) constitutes an adaptive drought-resistance mechanism remains debated. Consistent with the hydraulic fuse hypothesis (Bucci et al. 2013; Wason et al. 2018; Losso et al. 2019), recent syntheses have shown that both leaf–stem and root–stem segmentation tend to increase with climatic aridity, suggesting that hydraulic compartmentalization may become increasingly advantageous under water limitation (Rimer et al. 2024; Tang et al. 2024). Vulnerability segmentation has also been linked to drought-induced mortality patterns across a range of ecosystems, further supporting a potential adaptive role (Zhang et al. 2019; Huang et al. 2024a). Our results provide partial support for this hypothesis, as significant segmentation between distal and more central, permanent organs was observed in *F. sylvatica*, *A. alba*, *A. platanoides, T. cordata and P. abies* (Fig. 2, 3; Table S1, S3). In *Q. robur,* even though significant segmentation was not detected, coarse roots exhibited relatively greater embolism resistance than stems and leaves (Fig. S8; Table S3), potentially indicating preferential maintenance of belowground hydraulic function during drought (Lloret et al. 2004). However, the relationship between segmentation and drought resistance was not straightforward. Several species exhibiting clear segmentation (for example *F. sylvatica* and *P. abies*) have nonetheless suffered substantial mortality and growth declines in Central Europe during recent drought events (Schuldt et al. 2020; Martínez del Castillo et al. 2022; Weigel et al. 2023), indicating that segmentation alone does not guarantee drought resilience under severe stress. Conversely, *P. sylvestris*, which showed only minimal segmentation in our study, has experienced widespread drought-induced declines in growth and vitality across Central Europe (Schuldt et al. 2020; Arend et al. 2021; Haberstroh & Werner, 2022; Bennett et al. 2023; Knapp et al. 2024; Gribbe et al. 2024). Similarly, *P. menziesii*, which also exhibited low segmentation, is generally considered more drought-tolerant than *P. abies*, likely due to deeper rooting and greater access to soil water (Hackmann et al. 2026; Schumann et al. 2024), underscoring that drought resistance can be driven by water acquisition strategies rather than hydraulic vulnerability alone. In *Q. robur*, the relatively high embolism resistance of coarse roots (though not statistically significant) may reflect an adaptive strategy related to the species’ deep root system, which has been shown to enhance drought resilience by enabling access to deeper soil moisture reserves (Kahmen et al. 2022). These contrasting responses suggest that drought resistance is apparently only weakly related to hydraulic segmentation.

It is likely that the importance of vulnerability segmentation for the development of drought resistance in trees depends on the interaction of segmentation with other physiological and structural traits such as embolism resistance, stomatal regulation stringency, residual water loss, hydraulic capacitance, rooting depth, plasticity in root water uptake, and hydraulic recovery potential (Volaire, 2018; Gessler et al. 2026; Volaire et al. 2026). The positive correlation between segmentation and aridity reported in recent meta-analyses (Rimer et al. 2024; Tang et al. 2024) suggests that segmentation may be favoured under increasing drought exposure.

### Study limitations

Several methodological constraints should be considered when interpreting our results. First, measurements were conducted on 1-to 3-year-old saplings, which exhibit limited hydraulic capacitance and do not display the behaviour of mature plants. Xylem water potentials were measured only in leaves and fine roots, but not in stems and coarse roots, during dehydration. While leaf water potential is commonly used as a proxy for whole-plant water status, hydraulic decoupling between organs particularly during drought progression can compromise the assumption of hydraulic equilibrium (Körner 2019). To avoid related bias, our analysis restricted embolism thresholds to organs measured directly (thus excluding stem and coarse roots). Moreover, the optical vulnerability method primarily detects embolism formation in peripheral xylem tissues and may underestimate embolism occurring deeper in tissues of larger stems and roots (Brodribb et al. 2017; Losso et al. 2025). Furthermore, measurements of minimum conductance and embolism resistance were conducted at different time points during the growing season (minimum conductance was measured in late September), potentially introducing temporal variability due to seasonal plasticity in xylem development and embolism resistance (Domec et al. 2006; Torres-Ruiz et al. 2016; Huang et al. 2024b; Ma et al. 2024). These factors may affect the accuracy of the inferred hydraulic trait relationships and should be considered in the interpretation of results.

Despite these limitations, our study provides one of the most comprehensive assessments of xylem vulnerability across the whole hydraulic pathway of temperate trees. By quantifying embolism resistance and dehydration timing in leaves, stems, coarse roots, and fine roots of the same individuals, our study directly evaluates segmentation across organs that are rarely studied simultaneously (Li et al. 2024; Harrison Day et al. 2024). In particular, the inclusion of belowground organs represents a major strength, as roots are frequently excluded from vulnerability studies despite their critical role in drought resistance and recovery (Skelton et al. 2017; Rodriguez-Dominguez et al. 2018; Losso et al. 2019; Li et al. 2024). Furthermore, linking conventional water potential-based vulnerability metrics to novel data of vulnerability dynamics (dehydration time) provides a more holistic perspective on plant drought responses and highlights dehydration timing as an important, yet often overlooked, component of hydraulic function.

## Conclusion

By incorporating both water potential-based and time-dependent measures of xylem vulnerability across four plant compartments, our results demonstrate that embolism resistance and dehydration timing are distinct, yet interdependent, components of drought resistance. Both hydraulic (water potential thresholds) and temporal vulnerability (dehydration time) varied substantially among organs and species, independent of the species’ isohydricity. In broadleaved trees, the observed relationship between delayed dehydration and minimum conductance suggests that residual water loss after stomatal closure critically shapes drought progression and thus embolism risk. Therefore, drought responses emerge from the interaction of embolism resistance, dehydration dynamics, and whole-plant hydraulic organization. Our approach of linking xylem vulnerability thresholds to the temporal development of embolism across multiple organs provides a more comprehensive framework for understanding plant drought responses and offers a basis for improving predictions of tree performance under global change-type droughts.

## Acknowledgements

We thank Regina Helbig for support in establishing the experiments and care of plants and Heinz Coners for multiple technical support during the measurements. We thank Christopher Lucani and Beatrice Harrison Day for their methodological support during the measurements. We also thank Mechthild Stange, Annegret Kellner and Irmgard Gerstmann for their support with the measurement of Gmin. **AI use statement:** This work utilized the Qwen 3-72B model, a large language model developed by Alibaba Group, for refinement of parts of the text. All text was critically reviewed and edited by the corresponding author to ensure accuracy and academic integrity.

## Author contributions

CL and SSP conceived the idea and designed the study; FG collected the cavicam data with the support of SSP; LVR, FG measured Gmin data; SSP analyzed the data; SSP, CL, and FG wrote the manuscript. All authors all commented on and approved the final version.

## Data and Materials Availability

The data that support the findings of this study will be provided upon a request to the corresponding author SSP.

## References

Allen CD, Macalady AK, Chenchouni H, Bachelet D, McDowell N, Vennetier M, Kitzberger T, Rigling A, Breshears DD, Hogg EH (Ted), et al. 2010. A global overview of drought and heat-induced tree mortality reveals emerging climate change risks for forests. Forest Ecology and Management 259: 660–684.

Anderegg WRL, Klein T, Bartlett M, Sack L, Pellegrini AFA, Choat B, Jansen S. 2016. Meta-analysis reveals that hydraulic traits explain cross-species patterns of drought-induced tree mortality across the globe. Proceedings of the National Academy of Sciences 113: 5024–5029.

Arend M, Link RM, Patthey R, Hoch G, Schuldt B, Kahmen A. 2021. Rapid hydraulic collapse as cause of drought-induced mortality in conifers. Proceedings of the National Academy of Sciences 118.

Arend M, Peters RL, Zahnd C, Ognjenovic M, Hoch G, Kahmen A. 2026. Early onset of stomatal closure confounds current interpretations and applications of iso-/anisohydry theory. New Phytologist 249: 2616–2622

Bailey IW. 1913. The Preservative Treatment of Wood—II. The Structure of the Pit Membranes in the Tracheids of Conifers and their Relation to the Penetration of Gases, Liquids, and Finely Divided Solids into Green and Seasoned Wood. Journal of Forestry 11: 12–20a.

Bates D, Mächler M, Bolker B, Walker S. 2014. Fitting Linear Mixed-Effects Models using lme4. arXiv:1406.5823 [stat].

Bennett AC, Rodrigues de Sousa T, Monteagudo-Mendoza A, Esquivel-Muelbert A, Morandi PS, Coelho de Souza F, Castro W, Duque LF, Flores Llampazo G, Manoel dos Santos R, et al. 2023. Sensitivity of South American tropical forests to an extreme climate anomaly. Nature Climate Change 13: 967–974.

Blackman CJ, Creek D, Maier C, Aspinwall MJ, Drake JE, Pfautsch S, O’Grady A, Delzon S, Medlyn BE, Tissue DT, et al. 2019. Drought response strategies and hydraulic traits contribute to mechanistic understanding of plant dry-down to hydraulic failure (F Meinzer, Ed.). Tree Physiology 39: 910–924.

Blackman CJ, Pfautsch S, Choat B, Delzon S, Gleason SM, Duursma RA. 2016. Toward an index of desiccation time to tree mortality under drought. Plant, Cell & Environment 39: 2342–2345.

Brodribb TJ. 2020. Learning from a century of droughts. Nature Ecology & Evolution 4: 1007–1008.

Brodribb TJ, Bienaimé D, Marmottant P. 2016. Revealing catastrophic failure of leaf networks under stress. Proceedings of the National Academy of Sciences 113: 4865–4869.

Brodribb TJ, Carriqui M, Delzon S, Lucani C. 2017. Optical Measurement of Stem Xylem Vulnerability. Plant Physiology 174: 2054–2061.

Brodribb TJ, Cochard H. 2009. Hydraulic Failure Defines the Recovery and Point of Death in Water-Stressed Conifers. Plant Physiology 149: 575–584.

Bucci SJ, Scholz FG, Peschiutta ML, Arias NS, Meinzer FC, Goldstein G. 2013. The stem xylem of Patagonian shrubs operates far from the point of catastrophic dysfunction and is additionally protected from drought-induced embolism by leaves and roots. Plant, Cell & Environment 36: 2163–2174.

Camarero JJ, Colangelo M, Valeriano C, Gazol A, González de Andrés E, Alonso-Forn D, Voltas J, Torres-Ruiz JM, Delzon S, Badel E, et al. 2026. Xylem Pit Anatomy and Minimum Leaf Conductance Drive Drought Mortality in Pinus pinaster. Plant, Cell & Environment 49: 279– 294.

Cardoso AA, Kane CN, Rimer IM, McAdam SAM. 2022. Seeing is believing: what visualising bubbles in the xylem has revealed about plant hydraulic function (S Shabala, Ed.). Functional Plant Biology 49: 759–772.

Challis A, Blackman C, Ahrens C, Medlyn B, Rymer P, Tissue D. 2022. Adaptive plasticity in plant traits increases time to hydraulic failure under drought in a foundation tree (J Martinez-Vilalta, Ed.). Tree Physiology 42: 708–721.

Choat B, Brodribb TJ, Brodersen CR, Duursma RA, López R, Medlyn BE. 2018. Triggers of tree mortality under drought. Nature 558: 531–539.

Choat B, Jansen S, Brodribb TJ, Cochard H, Delzon S, Bhaskar R, Bucci SJ, Feild TS, Gleason SM, Hacke UG, et al. 2012. Global convergence in the vulnerability of forests to drought. Nature 491: 752–755.

Cochard H, Damour G, Bodet C, Tharwat I, Poirier M, Améglio T. 2005. Evaluation of a new centrifuge technique for rapid generation of xylem vulnerability curves. Physiologia Plantarum 124: 410– 418.

Copie A, Scotti-Saintagne C, Lefèvre F, Cochard H, Delzon S, Dumas P-J, Gounelle D, Herbette S, Jean F, Mariotte N, et al. 2025. Beyond Proxies: Towards ecophysiological indicators of drought resistance for forest management. Tree Physiology 49: tpaf090.

Creek D, Lamarque LJ, Torres-Ruiz JM, Parise C, Burlett R, Tissue DT, Delzon S. 2020. Xylem embolism in leaves does not occur with open stomata: evidence from direct observations using the optical visualization technique. Journal of Experimental Botany 71: 1151–1159.

Cuneo IF, Knipfer T, Brodersen CR, McElrone AJ. 2016. Mechanical Failure of Fine Root Cortical Cells Initiates Plant Hydraulic Decline during Drought. Plant Physiology 172: 1669–1678.

Domec J-C, Lachenbruch B, Meinzer FC. 2006. Bordered pit structure and function determine spatial patterns of air-seeding thresholds in xylem of Douglas-fir (*Pseudotsuga menziesii*; Pinaceae) trees. American Journal of Botany 93: 1588–1600.

Duursma RA, Blackman CJ, Lopéz R, Martin-StPaul NK, Cochard H, Medlyn BE. 2019. On the minimum leaf conductance: its role in models of plant water use, and ecological and environmental controls. New Phytologist 221: 693–705.

Gessler A, Grünzweig JM, Bigio L, Hartmann H, McDowell N, Krumm F, Bose AK, Rigling A, Bugmann H, Vitali V, et al. 2026. Shaping future forests: how can ecophysiology support climate-smart forest management? New Phytologist 250: 2778–2813.

Gohel D, Skintzos P. 2023. flextable: Functions for tabular reporting.

Gribbe S, Enderle L, Weigel R, Hertel D, Leuschner C, Muffler L. 2024. Recent growth decline and shifts in climatic growth constraints suggest climate vulnerability of beech, Douglas fir, pine and oak in Northern Germany. Forest Ecology and Management 566: 122022.

Haberstroh S, Werner C. 2022. The role of species interactions for forest resilience to drought. Plant Biology 24: 1098–1107.

Hackmann CA, Paligi SS, Mund M, Hölscher D, Leuschner C, Pietig K, Ammer C. 2026. Root water uptake depth in temperate forest trees: species-specific patterns shaped by neighbourhood and environment. Plant Biology 28: 872–887.

Haines FM. 1935. Observations on the Occurrence of Air in Conducting Tracts. Annals of Botany os-49: 367–379.

Hammond WM, Williams AP, Abatzoglou JT, Adams HD, Klein T, López R, Sáenz-Romero C, Hartmann H, Breshears DD, Allen CD. 2022. Global field observations of tree die-off reveal hotter-drought fingerprint for Earth’s forests. Nature Communications 13: 1761.

Hammond WM, Yu K, Wilson LA, Will RE, Anderegg WRL, Adams HD. 2019. Dead or dying? Quantifying the point of no return from hydraulic failure in drought-induced tree mortality. New Phytologist 223: 1834–1843.

Harrison Day BL, Brodersen CR, Brodribb TJ. 2024. Weak link or strong foundation? Vulnerability of fine root networks and stems to xylem embolism. New Phytologist 244: 1288–1302.

Harrison Day BL, Brodribb TJ, Walker NC, Brodersen CR. 2026. Fine Root Shrinkage Across Intact Root Networks Begins Early During Dehydration in Diverse Plants From Conifers to Angiosperms. Plant, Cell & Environment 49: 746–759.

Hartmann H, Link RM, Schuldt B. 2021. A whole-plant perspective of isohydry: stem-level support for leaf-level plant water regulation. Tree Physiology: tpab011.

Haverroth EJ, Rimer IM, Oliveira LA, de Lima LGA, Cesarino I, Martins SCV, McAdam SAM, Cardoso AA. 2024. Gradients in embolism resistance within stems driven by secondary growth in herbs. Plant, Cell & Environment 47: 2986–2998.

Hochberg U, Windt CW, Ponomarenko A, Zhang Y-J, Gersony J, Rockwell FE, Holbrook NM. 2017. Stomatal Closure, Basal Leaf Embolism, and Shedding Protect the Hydraulic Integrity of Grape Stems. Plant Physiology 174: 764–775.

Holbrook NM, Sinclair TR. 1992. Water balance in the arborescent palm, Sabal palmetto. II. Transpiration and stem water storage. *Plant*, Cell & Environment 15: 401–409.

Huang R, Wang Y, Duan J, Xi B, Yang J, Li X, Feng J, Choat B, Tissue DT. 2024a. Robust hydraulic traits correlation in woody species despite large trait variation along natural and experimental environmental gradients. Functional Ecology 38: 1835–1847.

Huang D-L, Zhang F, Qie Y-D, Wang Y-Q, Wan C-Y, Yu J-R, Zhang J, Gan J-X, Zhu S-D. 2024b. Vulnerability segmentation is vital to hydraulic strategy of tropical–subtropical woody plants. Journal of Ecology 112: 2798–2812.

Huo C, Villar-Salvador P, Li Y, Wang J, Li G. 2024. Physiological recovery after drought increases with leaf and taproot drought tolerance among oak seedlings. Environmental and Experimental Botany 222: 105747.

Jansen S, Kaack L, Ahmed MA, Beikircher B, Bittencourt P, Chen R, Flexas J, Gleason SM, Guha A, Heuret P, et al. 2026. Beyond air-seeding: Dynamic, multiphase interactions reveal a two-step mechanism of embolism propagation in angiosperm xylem. : 2026.08.05.743004.

Johnson DM, Domec J-C, Berry ZC, Schwantes AM, McCulloh KA, Woodruff DR, Polley HW, Wortemann R, Swenson JJ, Mackay DS, et al. 2018. Co-occurring woody species have diverse hydraulic strategies and mortality rates during an extreme drought. Plant, Cell & Environment 41: 576–588.

Johnson DM, Wortemann R, McCulloh KA, Jordan-Meille L, Ward E, Warren JM, Palmroth S, Domec J-C. 2016. A test of the hydraulic vulnerability segmentation hypothesis in angiosperm and conifer tree species. Tree Physiology 36: 983–993.

Kahmen A, Basler D, Hoch G, Link RM, Schuldt B, Zahnd C, Arend M. 2022. Root water uptake depth determines the hydraulic vulnerability of temperate European tree species during the extreme 2018 drought. Plant Biology 24: 1224–1239.

Kerstiens G. 1996. Cuticular water permeability and its physiological significance. Journal of Experimental Botany 47: 1813–1832.

Knapp N, Wellbrock N, Bielefeldt J, Dühnelt P, Hentschel R, Bolte A. 2024. From single trees to country-wide maps: Modeling mortality rates in Germany based on the Crown Condition Survey. Forest Ecology and Management 568: 122081.

Körner C. 2019. No need for pipes when the well is dry—a comment on hydraulic failure in trees. Tree Physiology 39: 695–700.

Kunz J, Räder A, Bauhus J. 2016. Effects of Drought and Rewetting on Growth and Gas Exchange of Minor European Broadleaved Tree Species. Forests 7: 239

Lens F, Gleason SM, Bortolami G, Brodersen C, Delzon S, Jansen S. 2022. Functional xylem characteristics associated with drought-induced embolism in angiosperms. New Phytologist 236: 2019–2036.

Lenth RV, Bolker B, Buerkner P, Giné-Vázquez I, Herve M, Jung M, Love J, Miguez F, Riebl H, Singmann H. 2023. emmeans: Estimated Marginal Means, aka Least-Squares Means.

Leuschner C. 2020. Drought response of European beech (*Fagus sylvatica* L.)—A review. Perspectives in Plant Ecology, Evolution and Systematics 47: 125576.

Leuschner C, Fuchs S, Wedde P, Rüther E, Schuldt B. 2024. A multi-criteria drought resistance assessment of temperate *Acer, Carpinus, Fraxinus, Quercus*, and *Tilia* species. Perspectives in Plant Ecology, Evolution and Systematics 62: 125777.

Leuschner C, Meinzer FC. 2024. Drought resistance and drought adaptation of Douglas-fir (*Pseudotsuga menziesii*) – A review. Perspectives in Plant Ecology, Evolution and Systematics 65: 125829.

Leuschner C, Wedde P, Luebbe T. 2019. The relation between pressure–volume curve traits and stomatal regulation of water potential in five temperate broadleaf tree species. Annals of Forest Science 76.

Li Y, Fan D, Huo C, Wang J, Wang X, Dong Z, Meng X, Li G. 2024. The degree of hydraulic vulnerability segmentation at the individual level is related to stomata sensitivity in seedlings of three oak species. Physiologia Plantarum 176: e14132.

Li R, Lu Y, Peters JM, Choat B, Lee AJ. 2020. Non-invasive measurement of leaf water content and pressure–volume curves using terahertz radiation. Scientific Reports 10.

Liang X, Cui B, Wang X, Liu H, He P, Yin D, Tang W, Wang Y, Liao L, Xia Q, et al. 2026. The coordination of dehydration tolerance and avoidance in oaks is mediated by leaf habit across a precipitation gradient. Functional Ecology 40: 750–763.

Lloret F, Siscart D, Dalmases C. 2004. Canopy recovery after drought dieback in holm-oak Mediterranean forests of Catalonia (NE Spain). Global Change Biology 10: 2092–2099.

Losso A, Bär A, Dämon B, Dullin C, Ganthaler A, Petruzzellis F, Savi T, Tromba G, Nardini A, Mayr S, et al. 2019. Insights from in vivo micro-CT analysis: testing the hydraulic vulnerability segmentation in *Acer pseudoplatanus* and *Fagus sylvatica* seedlings. New Phytologist 221: 1831–1842.

Lübbe T, Lamarque LJ, Delzon S, Torres Ruiz JM, Burlett R, Leuschner C, Schuldt B. 2022. High variation in hydraulic efficiency but not xylem safety between roots and branches in four temperate broad-leaved tree species. Functional Ecology 36: 699–712.

Lüdecke D, Ben-Shachar MS, Patil I, Waggoner P, Makowski D. 2021. performance: An R package for assessment, comparison and testing of statistical models. Journal of Open Source Software 6: 3139.

Ma B-L, Liao S-H, Lv Q-Z, Huang X, Jiang Z-M, Cai J. 2024. Seasonal plasticity of stem embolism resistance and its potential driving factors in six temperate woody species. Physiologia Plantarum 176: e14421.

Magh R-K, Eiferle C, Burzlaff T, Dannenmann M, Rennenberg H, Dubbert M. 2020. Competition for water rather than facilitation in mixed beech-fir forests after drying-wetting cycle. Journal of Hydrology 587: 124944.

Mantova M, Menezes-Silva PE, Badel E, Cochard H, Torres-Ruiz JM. 2021. The interplay of hydraulic failure and cell vitality explains tree capacity to recover from drought. Physiologia Plantarum 172: 247–257.

Martinez del Castillo E, Zang CS, Buras A, Hacket-Pain A, Esper J, Serrano-Notivoli R, Hartl C, Weigel R, Klesse S, Resco de Dios V, et al. 2022. Climate-change-driven growth decline of European beech forests. Communications Biology 5: 1–9.

Martínez-Vilalta J, Garcia-Forner N. 2017. Water potential regulation, stomatal behaviour and hydraulic transport under drought: deconstructing the iso/anisohydric concept. Plant, Cell & Environment 40: 962–976.

Martínez-Vilalta J, Poyatos R, Aguadé D, Retana J, Mencuccini M. 2014. A new look at water transport regulation in plants. New Phytologist 204: 105–115.

Martin-StPaul N, Delzon S, Cochard H. 2017. Plant resistance to drought depends on timely stomatal closure. Ecology Letters 20: 1437–1447.

McCulloh KA, Johnson DM, Meinzer FC, Woodruff DR. 2014. The dynamic pipeline: hydraulic capacitance and xylem hydraulic safety in four tall conifer species. Plant, Cell & Environment 37: 1171–1183.

McDowell N, Pockman WT, Allen CD, Breshears DD, Cobb N, Kolb T, Plaut J, Sperry J, West A, Williams DG, et al. 2008. Mechanisms of Plant Survival and Mortality during Drought: Why Do Some Plants Survive while Others Succumb to Drought? The New Phytologist 178: 719– 739.

Meinzer FC, James SA, Goldstein G, Woodruff D. 2003. Whole-tree water transport scales with sapwood capacitance in tropical forest canopy trees. Plant, Cell & Environment 26: 1147–1155.

Meinzer FC, Woodruff DR, Domec J-C, Goldstein G, Campanello PI, Gatti MG, Villalobos-Vega R. 2008. Coordination of leaf and stem water transport properties in tropical forest trees. Oecologia 156: 31–41.

Meinzer FC, Woodruff DR, Marias DE, Smith DD, McCulloh KA, Howard AR, Magedman AL. 2016. Mapping ‘hydroscapes’ along the iso-to anisohydric continuum of stomatal regulation of plant water status. Ecology Letters 19: 1343–1352.

Paligi SS, Hackmann CA, Schick J, Audisio M, Coners H, Mund M, Ammer C, Leuschner C. 2026. Isohydric Douglas fir shows earlier drought-induced declines in stem water status and growth than anisohydric beech. Agricultural and Forest Meteorology 388: 111294.

Paligi SS, Lichter J, Kotowska M, Schwutke RL, Audisio M, Mrak K, Penanhoat A, Schuldt B, Hertel D, Leuschner C. 2024. Water status dynamics and drought tolerance of juvenile European beech, Douglas fir and Norway spruce trees as dependent on neighborhood and nitrogen supply. Tree Physiology 44: tpae044.

Peters RL, Arend M, Zahnd C, Hoch G, Arndt SK, Cernusak LA, Poyatos R, Zhorzel T, Kahmen A. 2025. Uniform regulation of stomatal closure across temperate tree species to sustain nocturnal turgor and growth. Nat. Plants 11, 725–730. 10.1038/s41477-025-01957-3

Peters JMR, Choat B. 2025. Out on a Limb: Testing the Hydraulic Vulnerability Segmentation Hypothesis in Trees Across Multiple Ecosystems. Plant, Cell & Environment 48: 2162–2177.

Peters JMR, Gauthey A, Lopez R, Carins-Murphy MR, Brodribb TJ, Choat B. 2020. Non-invasive imaging reveals convergence in root and stem vulnerability to cavitation across five tree species. Journal of Experimental Botany 71: 6623–6637.

Pittermann J, Choat B, Jansen S, Stuart SA, Lynn L, Dawson TE. 2010. The Relationships between Xylem Safety and Hydraulic Efficiency in the Cupressaceae: The Evolution of Pit Membrane Form and Function. Plant Physiology 153: 1919–1931.

Pörtner H-O, Roberts DC, Tignor MMB, Poloczanska ES, Mintenbeck K, Alegría A, Craig M, Langsdorf S, Löschke S, Möller V, et al. (Eds). 2022. Climate Change 2022: Impacts, Adaptation and Vulnerability. Contribution of Working Group II to the Sixth Assessment Report of the Intergovernmental Panel on Climate Change.

Preisler Y, Hölttä T, Grünzweig JM, Oz I, Tatarinov F, Ruehr NK, Rotenberg E, Yakir D. 2022. The importance of tree internal water storage under drought conditions. Tree Physiology 42: 771– 783.

Pya N. 2025. scam: Shape constrained additive models. R package version 1.2-20, https://CRAN.R-project.org/package=scam

R Core Team. 2022. R: A language and environment for statistical computing. Vienna, Austria.

Rimer I, Kane C, Yan D, Paudel I, Dukes J, McAdam S. 2024. Considerable variation in embolism resistance occurs within plant communities and differentiates plant biomes. Preprint.

Rodriguez-Dominguez CM, Carins Murphy MR, Lucani C, Brodribb TJ. 2018. Mapping xylem failure in disparate organs of whole plants reveals extreme resistance in olive roots. New Phytologist 218: 1025–1035.

Sack L, Cowan PD, Jaikumar N, Holbrook NM. 2003. The ‘hydrology’ of leaves: co-ordination of structure and function in temperate woody species. *Plant*, Cell & Environment 26: 1343–1356.

Schoch W, Heller I, Schweingruber FH, Kienast F. (2004). Wood anatomy of central European Species. Available online at www.woodanatomy.ch.

Scholz FG, Bucci SJ, Goldstein G, Meinzer FC, Franco AC, Miralles-Wilhelm F. 2007. Biophysical properties and functional significance of stem water storage tissues in Neotropical savanna trees. Plant, Cell & Environment 30: 236–248.

Schuldt B, Buras A, Arend M, Vitasse Y, Beierkuhnlein C, Damm A, Gharun M, Grams TEE, Hauck M, Hajek P, et al. 2020. A first assessment of the impact of the extreme 2018 summer drought on Central European forests. Basic and Applied Ecology 45: 86–103.

Schumann K, Schuldt B, Fischer M, Ammer C, Leuschner C. 2024. Xylem safety in relation to the stringency of plant water potential regulation of European beech, Norway spruce, and Douglas-fir trees during severe drought. Trees. 38: 607–623.

Song Y, Sterck F, Zhou X, Liu Q, Kruijt B, Poorter L. 2022. Drought resilience of conifer species is driven by leaf lifespan but not by hydraulic traits. New Phytologist 235: 978–992.

Sorek Y, Greenstein S, Hochberg U. 2022. Seasonal adjustment of leaf embolism resistance and its importance for hydraulic safety in deciduous trees. Physiologia Plantarum 174: e13785.

Sperry JS, Tyree MT. 1988. Mechanism of Water Stress-Induced Xylem Embolism. Plant Physiology 88: 581–587.

Tang W, Liu X, Liang X, Liu H, Yu K, He P, McAdam S, Zhao H, Ye Q. 2024. Hydraulic vulnerability difference between branches and roots increases with environmental aridity. Oecologia 205: 177–190.

Torres-Ruiz JM, Cochard H, Mencuccini M, Delzon S, Badel E. 2016. Direct observation and modelling of embolism spread between xylem conduits: a case study in Scots pine. Plant, Cell & Environment 39: 2774–2785.

Trueba S, Burlett R, Bouteiller X, Burneau T, Forget G, Larter M, N’Do D, Ziegler C, Delzon S. 2026. Ecological drivers and phylogenetic patterns of leaf minimum conductance variability in vascular plants. New Phytologist 250: 3716–3731.

Tyree MT, Ewers FW. 1991. The hydraulic architecture of trees and other woody plants. New Phytologist 119: 345–360.

Tyree MT. 2003. Plant hydraulics: The ascent of water. Nature 423: 923–923.

Urli M, Porte AJ, Cochard H, Guengant Y, Burlett R, Delzon S. 2013. Xylem embolism threshold for catastrophic hydraulic failure in angiosperm trees. Tree Physiology 33: 672–683.

Volaire F. 2018. A unified framework of plant adaptive strategies to drought: Crossing scales and disciplines. Global Change Biology 24: 2929–2938.

Volaire F, Gleason SM, Carminati A, Griffin-Nolan RJ, Mencuccini M, Barkaoui K, Bucci S, Carriquí M, Comas L, Coupel-Ledru A, et al. 2026. From growth potential to drought survival: a trait- and time-based framework for plant water economics across vascular species. New Phytologist 250: 21–40.

Waite P-A, Kumar M, Link RM, Schuldt B. 2024. Coordinated hydraulic traits influence the two phases of time to hydraulic failure in five temperate tree species differing in stomatal stringency. Tree Physiology 44: tpae038.

Waite P-A, Riedel VP, Link RM, Beldman I, Kumar M, Nain P, Panagoulias M, Schuldt B. 2026. Coordinated hydraulic traits predict the time to critical hydraulic failure in temperate broad-leaved and conifer tree species. Plant Physiology 201: kiag177.

Wason JW, Anstreicher KS, Stephansky N, Huggett BA, Brodersen CR. 2018. Hydraulic safety margins and air-seeding thresholds in roots, trunks, branches and petioles of four northern hardwood trees. New Phytologist 219: 77–88.

Weigel R, Bat-Enerel B, Dulamsuren C, Muffler L, Weithmann G, Leuschner C. 2023. Summer drought exposure, stand structure, and soil properties jointly control the growth of European beech along a steep precipitation gradient in northern Germany. Global Change Biology 29: 763–779.

Wickham H. 2016. ggplot2: elegant graphics for data analysis. Springer-Verlag New York.

Wickham H, François R, Henry L, Müller K, RStudio. 2019. dplyr: A Grammar of Data Manipulation.

Wilkening JV, Skelton RP, Feng X, Dawson TE, Thompson SE. 2023. Exploring within-plant hydraulic trait variation: A test of the vulnerability segmentation hypothesis. Plant, Cell & Environment 46: 2726–2746.

Wu M, Zhang Y, Oya T, Marcati CR, Pereira L, Jansen S. 2020. Root xylem in three woody angiosperm species is not more vulnerable to embolism than stem xylem. Plant and Soil 450: 479-495.

Zambonini D, Savi T, Rosner S, Petit G. 2024. Consistent decrease in conifer embolism resistance from the stem apex to base resulting from axial trends in tracheid and pit traits. Frontiers in Plant Science 15.

Zeppan JK, Hülsmann L, Knüver T, Ziegler Y, Ruehr NK. 2026. Water Availability and Hydraulic Strategies Control Leaf Thermoregulation and Damage During Heat Stress and Recovery in Temperate Tree Species. Plant Cell & Environment 49: 6296–6310.

Zhang S, Wen G, Yang D. 2019. Drought-Induced Mortality Is Related to Hydraulic Vulnerability Segmentation of Tree Species in a Savanna Ecosystem. Forests 10: 697.

Zimmermann MH. 1983. Xylem Structure and the Ascent of Sap. Berlin, Heidelberg: Springer.

